# A Stochastic Dynamic Operator framework that improves the precision of analysis and prediction relative to the classical spike-triggered average method, extending the toolkit

**DOI:** 10.1101/2024.05.10.593606

**Authors:** Trevor S. Smith, Maryam Abolfath-Beygi, Terence D. Sanger, Simon F. Giszter

## Abstract

Here we test the Stochastic Dynamic Operator (SDO) as a new framework for describing physiological signal dynamics relative to spiking or stimulus events. The SDO is a natural extension of existing spike-triggered averaging (STA), or stimulus-triggered averaging, methods currently used in neural analysis. It extends the classic STA to cover state-dependent and probabilistic responses where STA may fail. SDO methods are more sensitive and specific than the STA for identifying state-dependent relationships in simulated data. We have tested SDO analysis for interactions between electrophysiological recordings of spinal interneurons, single motor units, and aggregate muscle electromyograms (EMG) of major muscles in the spinal frog hindlimb. When predicting target signal behavior relative to spiking events, the SDO framework outperformed or matched classical spike-triggered averaging methods. SDO analysis permits more complicated spike-signal relationships to be captured, analyzed, and interpreted visually and intuitively. SDO methods can be applied at different scales of interest where spike-triggered averaging methods are currently employed, and beyond, from single neurons to gross motor behaviors. SDOs may be readily generated and analyzed using the provided *SDO Analysis Toolkit*. We anticipate this method will be broadly useful for describing dynamical signal behavior and uncovering state-dependent relationships of stochastic signals relative to discrete event times.

**SIGNIFICANCE:** Here the authors introduce new tools and demonstrate data analysis using a new probabilistic and state-dependent technique, which is an expansion and extension of the classical spike-triggered average, the Stochastic Dynamic Operator. Stochastic Dynamic Operator methods extend application into domains where classical spike triggered averages fail, capture more information on spike correlations, and match or outperform the spike-triggered average when generating predictions of signal amplitude based on spiking events. A data and code package toolkit for utilizing and interpreting Stochastic Dynamic Operator methods is provided together with example analyses. Both the method and the associated toolkit are thus expected to be broadly useful in research domains where the spike triggered average is currently used for analysis, and beyond.

## INTRODUCTION

Standard methods of examining the association of a neural spike and an external or physiological signal, despite variability in environment and organismal state, involve averaging peri-event segments of the signal over multiple spiking events (i.e., ‘Spike-triggered’ averaging, STA; Kirkwood and Sears, 1975). Under various assumptions (e.g., noise is additive and independent, the spike effect is stationary), such averaging reduces the magnitude of stochastic noise, and isolates the correlation of a particular spike train and signal relative to other neurons or neural signals. This relationship between a spike and signal may vary: Spike may be affected by signal, spike may ‘effect’ a change of the signal, or a neuron may be both sensitive to and evoke a change in signal. Coarsely, analysis of individual neurons often seeks to characterize *when* a neuron is active, and *what happens* when the neuron fires.

Consider the role of the STA in predicting a change in signal amplitude. Spike-triggered averaging has identified significant structural and functional relationships from cortical neurons (e.g., Fetz and Cheney, 1980; Buys, Lemon, Mantel, and Muir, 1986), midbrain (e.g., Cheney, 1980), and spinal interneurons (e.g., Maier, Perlmutter, and Fetz, 1998; Hart and Giszter, 2010; Takei et. al., 2017) to muscle activations. STA has also been instrumental in characterizing receptive fields of neurons within the sensory systems. The related *stimulus-* triggered averaging has similarly been valuable for interpreting behavioral output responses (e.g., Cheney and Fetz, 1985) and is standard procedure for neural analysis. Importantly, triggered averaging necessarily presumes that signal behavior and evoked spike/stimulus effects are comparable across all spiking events (and hence can be averaged together). In practice, such relationships are often dynamic, such as the current length of a muscle affecting the tension generated by a motoneuron spike (Gordon, et al, 1966).

However, even under similar experimental recording conditions, neurons may demonstrate variable post-spike effects due to underlying neural system state or dynamics. To illustrate, consider the simplified spinal reflex pathway with monosynaptic spindle and disynaptic Golgi tendon organ (GTO) effects (**Figure 1A**). Here, some of the mechanical parameters of the muscle are encoded by muscle spindles and GTOs and relayed to the spinal cord via the Ia and Ib afferent, respectively. The Ia afferent in turn excites motoneurons within the homonymous motor pool monosynaptically, evoking muscle contraction in a textbook fashion (**Fig. 1B**). Spikes recorded at the Ia afferent are predictive of motor outflow, as measured by electroneurogram (ENG) or electromyogram (EMG).

**Figure 1:**
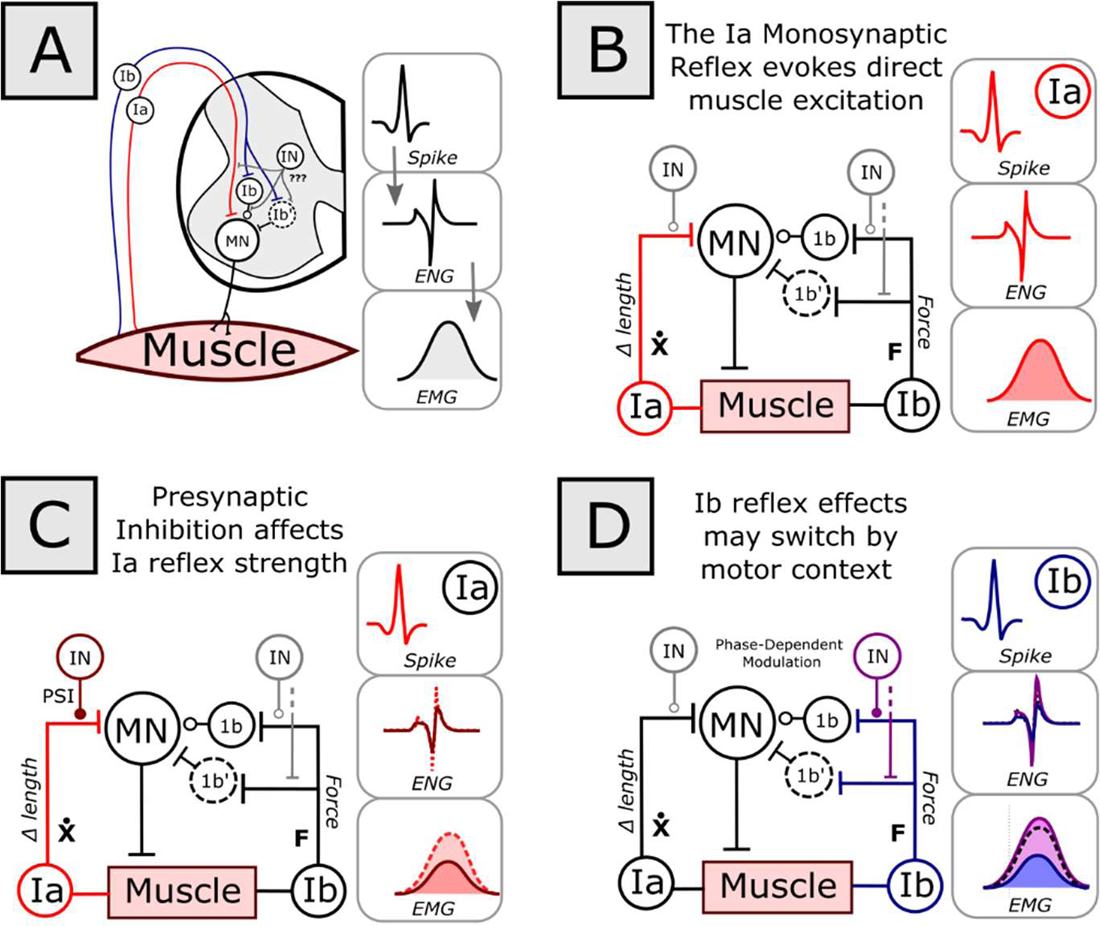
The nervous system generates state-dependent behavior. **A)** Due to the presence of monosynaptic and disynaptic reflexes provided by the Ia and 1b afferent systems, there is a parametric relationship between activity of these neurons and muscle behavior. One method of interrogating the effects of a neuron’s spike on the spinal output would be to average the electroneurogram (ENG) or electromyogram (EMG) signal traces at time of spike (i.e., ‘Spike-triggered Average’). Here, the state of other spinal interneurons which may influence this circuit are often not directly known. **B**) Within a simplified circuit model, through the monosynaptic reflex, the Ia afferent detects changes in muscle length and provides excitation to the homonymous motor pool. Under these conditions, the STA effect of the IA afferent will be to increase the ENG and EMG of this muscle. However, the Ia Afferent is only one of many inputs into the system. **C)** Presynaptic inhibition (PSI) of the Ia afferent may directly attenuate the amplitude of the Ia afferent STA. If this average Trigger averaging EMG relative to the spikes of the Ia afferent will necessarily average over both conditions, which may obscure dynamical behavior. **D**) The effects of the Ib afferents are known to be selectively modulated by spinal state/activity. While in some conditions the Ib afferent may attenuate ENG or EMG levels to the homonymous motor pool (blue), in other conditions, the Ib afferent spike-effects may enhance this activity (purple). Clearly, using a simple spike-triggered average across these paradoxical conditions would be ill-equipped to handle this behavior. MN = Motoneuron, IN = Interneuron, PSI = Presynaptic Inhibition.

(Indeed, direct stimulation of this pathway induces the Hoffmann reflex (Hoffmann, 1910), comprising a clinical measure of spinal excitability.) However, the precise parametric relationship between Ia afferent and muscle response also depends on other factors, e.g., muscle length (Shiavi and Negin, 1975) and presence of presynaptic inhibition of the Ia afferent in the spinal cord (Devanandan, Eccles, and Yokota, 1964)(**Fig. 1B**). Presynaptic inhibition (as modelled by IN in **Fig. 1C**), or other inhibitory drives may cause Ia afferent spikes to fail to evoke excitatory postsynaptic potentials on motoneurons or lessen observed ENG or EMG responses. The effective strengths of Ia reflexes are known to be actively regulated, and to differ among spinal conditions, such as posture and locomotion, and between locomotor phases. The correlation between a spike of the Ia afferent (itself only one of many converging inputs to the motor pool) and motor behavior may thus depend upon the state of these ongoing background spinal dynamics. Indeed, the Ib pathway’s state-driven modulation is still more influential and may switch inhibition to excitation based on locomotor phase (**Fig. 1D**)(Prochazka, 1997). Aspects of these background dynamics are also reflected in *signal* ‘state’ (here, the level of ENG or EMG activity). Spike-triggered averaging of the effect of the Ia or Ib afferent, without considering signal state, also averages over these intrinsic dynamics and potential state-dependent modulations.

Our objective here is to present a flexible and scalable data modelling and analysis framework which can better capture dynamical signal behaviors in the central nervous system relative to spiking events, where state dependence exists, and where responses may be stochastic. Previously, the ‘Stochastic Dynamic Operator’ (SDO) was developed and demonstrated as a theoretical model accounting for stochastic state-dependent effects of individual and population neural firing on signal dynamics (Sanger, 2010, 2011, 2014). Here, for the first time, we demonstrate the utility of SDOs for analysis and interpretation of real neural datasets. We briefly explore examples of SDO analysis for interpreting and stochastically predicting biological signals using the correlated activity between interneurons, single motor units, and muscles in spinal motor behaviors.

## MATERIALS AND METHODS

### Humane treatment and Institutional Animal Care and Use Committee approvals

All experimental procedures complied with the guidelines of the National Institutes of Health Guide for Care and Use of Laboratory Animals and received approval from the Institutional Animal Care and Use Committee of Drexel University.

### Biological Rationale and Choice of Test Data Framework

We selected the well-worked and robust spinal frog model to test the SDO framework with biological data. The spinal frog exhibits a well-characterized hindlimb-to-hindlimb wiping reflex (Giszter et al, 1989; Schotland and Rymer, 1993a; Schotland and Rymer, 1993b; Kargo and Giszter, 2000) to rid the skin of a noxious stimulus. The role of spinal interneurons contributing to muscle activations within this model has previously been investigated using STA (Hart and Giszter, 2010). Furthermore, because the frog lacks gamma motor neurons, proprioceptive feedback from the musculature is dependent upon muscle activations, lengths, and forces, which may be experimentally measured or estimated, and is not modified by independent control of muscle spindles. This anatomical arrangement is ideal for minimizing trial-to-trial variation in reflexive wiping, representing an ideal case for the use of classical STA within the system, and to compare STA in the best light with the new SDO approaches.

### Surgical Methods

We anesthetized adult bullfrogs using 1 ml/ kg of 5% tricaine and incubated them on ice to quicken the anesthetic effect. The skin on the skull was incised along the dorsal midline between the eyes and ears, then reflected away to reveal the musculature. We incised the midline musculature fascia behind the skull and deflected the musculature using a retractor to expose the foramen magnum. After opening the foramen, we cauterized the vascular dura over the fourth ventricle and cut an opening along the midline with iridectomy scissors. We then laterally transected the spinal cord with fine vacuum aspiration immediately below the medulla of the frog, taking care not to rupture any of the remaining dura or large blood vessels along the sides of the incision or underlying the spinal tissue. Following spinal transection, we filled the cavity with a piece of Gelfoam and closed the incision. We next made a small opening over the brain and decerebrated the frog at a low level through repeated applications of heat cautery to the tectum and rostral structures. After packing the cavity with Gelfoam, the incision was closed with wound clips and sealed with Vetbond (cyanoacrylate). We inserted bipolar intramuscular stainless steel electromyography (EMG) electrodes in 11 muscles of the frog right hindlimb: the rectus anterior (RA), rectus interior (RI), adductor magnus (AD), semimembranosus (SM), gluteus (GL), vastus internus (VI), biceps (BI), sartorius (SA), vastus externus (VE), gastrocnemius (G), and tibialis anterior (TA). For recording from the fragile sartorius (SA), a silicon patch electrode was placed under the muscle belly. To expose the spinal cord for extracellular recording, we made an incision on the mid-to-lower back of the frog’s skin and musculature and reflected the skin back. We incised fascia, separated the back musculature with blunt dissection, and kept the muscles deflected via retractors. Other vertebral musculature was cleared using blunt dissection and iridectomy scissors until the bone of the vertebral arches was cleanly exposed. After clearing connective tissue, we used rongeurs to cut away the spinal arch and spinous processes, revealing the dura beneath. We removed three arches, exposing the L2/L3 border region of the frog spinal cord. We used an electrocautery to cauterize small patches of blood vessels in the vascular dura and expanded these holes until it was possible to deflect the dura and the attached blood vessel back, revealing the spinal cord pia mater for a length of three spinal segments. This allowed full access to half the width of the spinal cord, from the midline to the right lateral extreme of the cord. We covered the dura with moistened Gelfoam and a cotton ball. The frog was allowed to recover overnight. We opened a small hole in the pia mater using microincision on the day of recording to provide access directly to the white and gray matter.

### Reflex Preparation

During experimental recordings, the frog was placed on a molded stand that supported the body in the horizontal plane. The pelvis and vertebral column were immobilized using custom-made clamps. The wiping limb was secured into an ankle restraint connected to a six-axis force/torque transducer (ATI 310) to record isometric forces generated at the ankle. Hindlimb-to-hindlimb wiping was evoked by electrically stimulating the dorsolateral surface of the heel of the target (left, contralateral) limb. The stimulation was delivered via bipolar leads (2–3 mm separation) and consisted of a 350ms train of 3–8 V, 1 ms biphasic pulses applied at 40 Hz. This stimulus evoked either bilaterally-coordinated hindlimb wiping attempts, between the contralateral limb (containing EMG electrodes) and ipsilateral limb, or ipsilateral flexion-withdrawals from the stimulated limb. We adjusted the bipolar leads as necessary to evoke wiping motor responses from the right (contralateral) limb. Trials were composed of 60 second intervals, with stimuli applied 5-10 seconds after trial recording. Stimulus applications were spaced at least 2 minutes apart to avoid habituation.

### EMG recording

EMG signals were recorded from the right, isometrically-fixed, hindlimb using implanted wire pairs. Signals were sampled at 2 kHz, differentially amplified by a gain of 10,000 by a Ripple™ Grapevine EMG front-end amplifier, and band-pass filtered (15-375 Hz) online by a Ripple™ Grapevine neural interface processor (**Fig 2A**). Spinally-organized wiping movements were elicited through a cutaneous stimulation electrode applying a 250 ms train of 40 Hz biphasic 1 ms electrical pulses to the ankle dorsum.

**Figure 2:**
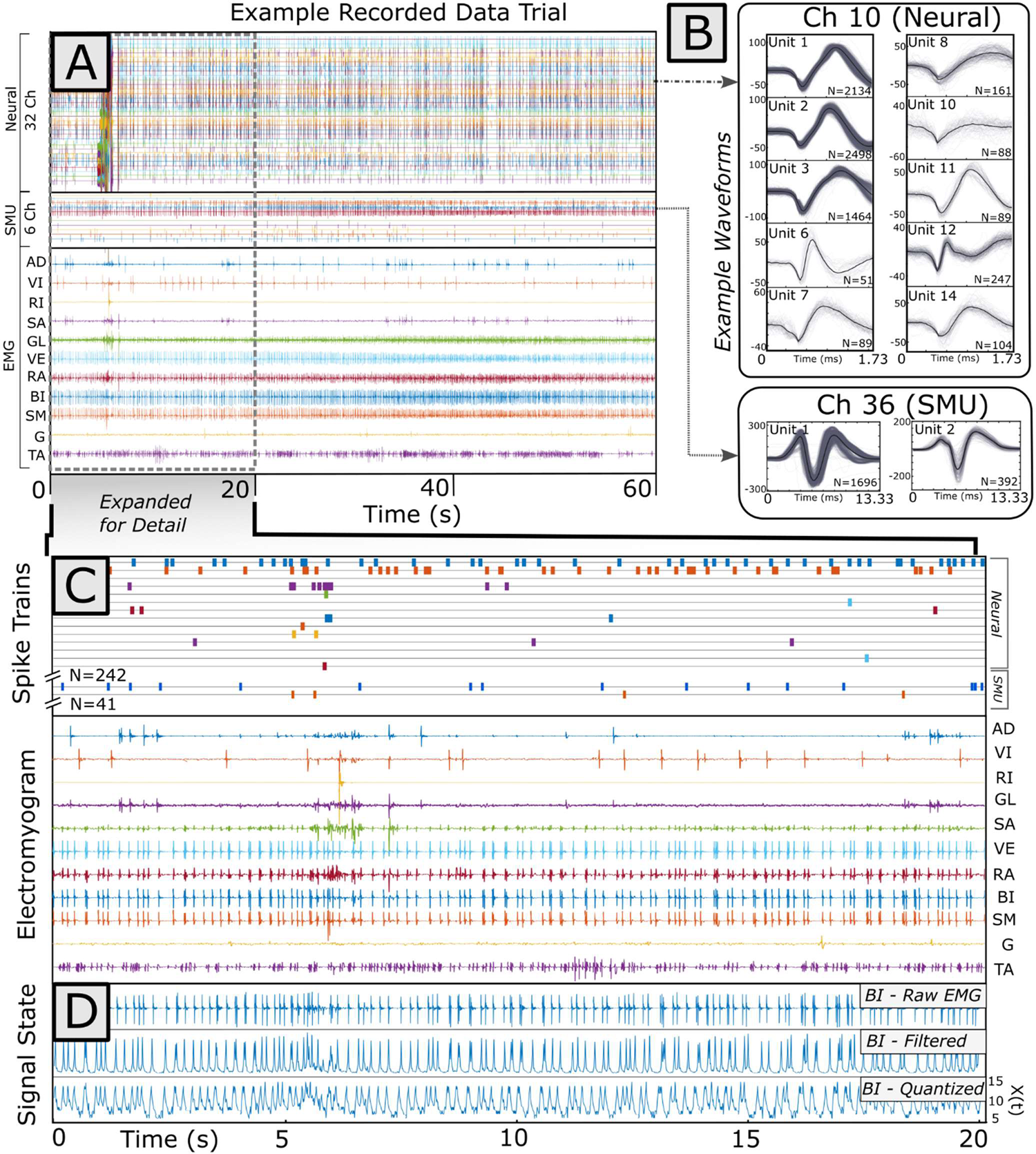
Example 60-second data trial. **A**) Spiking data were recorded on 32 channels (neurons) and 6 data channels (motor units). Analog EMG was recorded across 11 muscles. Both sensory-elicited and ongoing motor behavior were recorded in this trial. **B**) Example waveforms from a single neural and motor unit channel within the greater dataset. Waveforms from both neural and motor units, collected across all trials, were sorted in *Plexon Offline Sorter*. (The 13ms SMU waveforms displayed here were extracted from stainless-steel vastus externus EMG signal recorded 3.33 ms before to 10 ms *after* SMU spike time.) Spiking events in sorted clusters were treated as point processes. **C**) Expanded signal and sorted point processes from a single neural and motor unit channel. The total number of spikes and their correlated occurrence relative to EMG signal could be investigated through different analytical methods, including the spike-triggered average or the stochastic dynamic operator. **D**) Example raw EMG from the biceps femoris muscle, which was rectified and filtered, then quantized to derive the time series of signal ‘state’ used for later analysis. Here higher signal ‘state’ corresponds to a higher signal amplitude of the rectified and filtered EMG envelope.

### Neural Recording

We recorded extracellular neural activity using a flexible braid (Kim et al, 2019A, Kim et al, 2019B). of 24 channel microwires (each wire 9.6 μm diameter) forming a multi-electrode, ∼240 μm total in diameter Recording site impedances on the microwire were set between 0.2 – 0.5 MΩ by controlled platinum nanograss plating. We targeted the L1-L3 region of the spinal cord, < 500 μm lateral of the spinal cord midline, a region of the spinal cord with a high-yield of motor responses to microstimulation and high-yield of wipe-active neurons (Bizzi et al, 1991; Giszter et al, 1993; Hart and Giszter, 2010). The neural multi-electrode was advanced until action potentials were apparent (**Fig. 2A**). Continuous signals were filtered using 0.3Hz High-pass, and 7.5 kHz Low-pass filters, and digitized at 30 kHz using a Ripple™ Grapevine neural interface using *Nano2* Front Ends. Neural events were detected using a threshold of 4.5x the root-mean square (RMS) of each channel’s background activity. Each event was sampled as a 1.73 ms (52-point) waveform for offline clustering and analysis.

### Motor Unit Recording

Motor unit action potentials were recorded in muscle using a pair of 6-channel braided microwire (9.6 μm diameter) electrodes, with a seventh wire acting as reference, also in the braid assembly. Microwires were of sufficient length to reach a Ripple™ Grapevine Neural Front-End Amplifier without tension during frog movement (∼8-10 cm). Motor unit data was recorded as with neural data, above. We exposed a single recording site (∼50 μm) on each 9.6 μm Nickel-Chromium wire in the braid using laser microablation, with site impedance controlled to 0.2 – 0.5 MΩ via gold electroplating, like the neural probe. Single motor unit probes were inserted into the proximal vastus externus muscle, oblique to the fiber pennation angle, via a curved suture needle. Electrodes were also grounded through a 7-strand stainless steel wire (A-M Microsystems) into the belly of the same muscle.

## DATA PROCESSING METHODS

### Data Preprocessing

Offline cluster cutting of both interneuron and motor unit spikes was performed using Plexon Neurotechnology Research System’s Offline Sorter software. The first three principal components of 1.73 ms-long waveforms, centered on each recorded spike, were clustered using an automated, expectation-maximization sorting algorithm (Figueiredo and Jain, 2002; Shoham et al., 2003) followed by manual curation (**Fig. 2B**). Sorted spike trains and EMG signals were imported into MATLAB for further analysis (**Fig. 2C**). EMG signals were further processed offline using a series of zero-phase filters: 1) a 60 Hz notch filter, 2) a 10 Hz high-pass filter (4^th^ order Butterworth), 3) and 20-point root-mean square (RMS) filter, which smoothed and rectified the EMG signal. We assigned EMG signal ‘state’ based on signal amplitude. Since the smoothed and rectified EMG signal amplitude distribution fell into a roughly exponential distribution, we quantized EMG amplitude into signal states using logarithmic binning to allocate probabilities of state closer to a normal distribution: We defined 20 signal states for each EMG channel, representing equal intervals of the log-transformed EMG signal amplitude (**Fig. 2D**).

### ALGORITHMS DEVELOPED AND TESTED

#### Theoretical Consideration of when the Spike-Triggered Average May Fail

Biological systems are dynamic and incompletely observed. Consequently, signals measured within the system are often stochastic, and spuriously correlated from an outside perspective. When determining a neurons’ contribution or sensitivity to a measured covariate signal, various assumptions are made regarding the structure of the relationship. The STA supports different causal models. STA provides one method to reverse-correlate a neuron’s spikes to preceding changes in signal amplitude. STA also provides a means to predict future signal state based on spike occurrence. The latter is the focus of the methos compared here. When spiking events (or at least the result of neural spikes) are considered independent and the stochastic component of the signal is uncorrelated with spike, averaging signal envelope around spiking events provides a straightforward method of isolating a single ‘response’ of the signal corresponding to spiking events.

The time series data used in the STA comprises signal amplitude, *x,* indexed at time *t*, i.e., x(t) (Generally, t is collected at the limit of digital signal sampling) (**Fig. 3A**). To mechanistically compute the STA, pre-spike and post-spike signal levels are sampled over some finite interval, *Δt*, relative to spike time, *s*. For every time point relative to spike, *t*_i_ ∈ (*s* − Δ*t, s* + Δ*t*), concurrent signal amplitude, *x(ti),* is averaged over all spiking events, to find the mean, x^-^(ti) (**Fig. 3C**). These signal waveforms may be variously pre-processed prior to this averaging operation (e.g., mean-leveling) to further isolate spike-associated correlations. Performing this operation on all relative time points thus generates a mean waveform, x^-^(t), which is effectively the signal impulse response to the spiking impulse.

**Figure 3:**
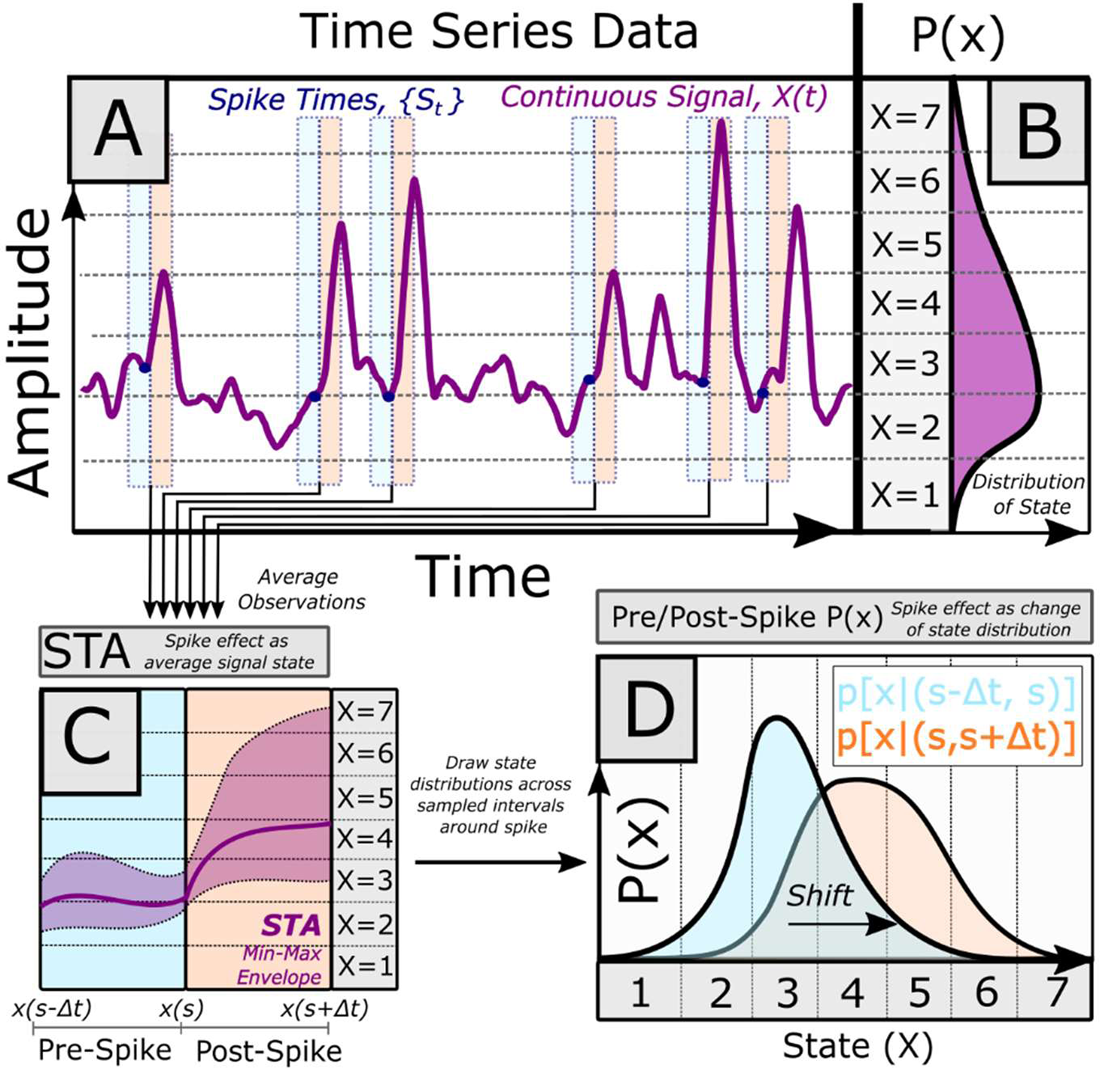
**A)** Electrophysiological recordings are often time series data: Time-varying amplitude values collected over a duration. The relationship of a point process, such as a spike train, and a time series is often evaluated as the spike-triggered average (STA). Here, intervals of the continuous signal are sampled around spiking events, with samples averaged together. **C**) Under the STA framework, if a spike-signal relationship is significant, an ‘impulse response’ may be present, showing a difference in the average signal behavior in the pre-spike vs. post-spike averaged interval. **B)** A complementary approach is to treat the electrophysiological signal as a ‘discrete random variable. By applying a quantization scheme, continuous signal amplitudes may be converted into a discrete time series of finite signal ‘states’. This permits signal behavior over an interval to then be described as a probability distribution. **D)** As with the classical spike-triggered average (**C**), the relationship between a spike and signal may be discerned from intervals of the discrete signal before and after spike. The analogous spike-triggered average probability distributions describe signal state prior to, and after, a spike. The correlated effect of a spike event on signal behavior can thus be described as the shift in the spike-triggered pre-spike and post-spike signal distributions.

In this framework, the neuron’s contribution to the correlated signal (or the signal’s effective stimuli, if signal facilitates spike in reverse correlation) is the convolution of the spiking impulses with the expected impulse response (i.e., the STA). Here, spikes are implicitly considered as independent and equivalent events, with the spike rate reflecting neural excitation (i.e., the neuron behaves within the assumptions of the Linear-Nonlinear Poisson model) (Schwartz, et al., 2006). If these assumptions are upheld, then the STA should provide the maximum likelihood estimation of the spike-signal correlation (Paninski, 2004).

These assumptions bear consideration. Is a neural spike equally potent in predicting future signal state across all conditions? An initial exploration of this question may be directed at the covariate signal itself: Does the spike-signal correlation depend on signal amplitude? It is important to recognize that the STA operation enforces a type of dimensionality reduction: For each position in time relative to spike, *t*, the population of observed signal responses form a probability distribution, *p(x,t)*, from which the classical STA merely utilizes the mean, a single value, thus reporting the ‘average’ response, and a gaussian (normal) distribution assumption for statistical significance. If the envelope is first mean-leveled prior to averaging, then any consideration of amplitude-dependent spike correlations is also lost. In estimating a single response, the STA thus contains less complete information about signal behavior at time *t* compared to the full distribution of signal values around the spike.

#### The Spike-Triggered Average Distribution of Signal States

We here propose a method of utilizing the complete spike-triggered state probability distribution to describe spike-triggered effects. Our process is as follows: We first apply an amplitude binning scheme to our recorded time series data (e.g., EMG), such that each observation of the recorded signal amplitude may be assigned to a single discrete amplitude bin, *xi.* (The number of such bins can be chosen for the signal based on signal range and precision required) (**Fig. 3A,B**). Here we define signal “state” as a binned range of signal amplitude, resultant of the quantization scheme applied to the original, continuous, time series data. (Hence, “higher states” corresponds to higher levels of original signal amplitude.) Here, the frequency of observing state over the entire time series is described by the distribution P(x) (**Fig. 3B**). Next, we infer the spike-triggered effect, classically inferred from the change in the amplitude (waveform) before and after spike as the change in the *distributions* of signal amplitude around each spike (**Fig. 3D**). As with the classical STA, signal is sampled over a short interval (s-*Δt, s+Δt*) relative to the spike event, *s* (**Fig. 4A**). By quantizing signal into discrete state, potentially non-normally distributed signal amplitudes in the STA (**Fig. 4B**) can be described explicitly for any time point as a distribution of state (**Fig. 4C**). We define the *pre-spike* distribution as the distribution of signal states over the entire sampled interval leading to spike *p*[*x*, (*s* − Δ*t, s*)], and the *post-spike* distribution as the distribution of states in the entire sampled interval after spike, *p*[*x*, (*s, s* + Δ*t*)] (**Fig. 4D**). (For brevity, we will represent the *pre-spike* and *post-spike* distributions as *p*(*x*_o_) and *p*(*x*_l_), respectively.) Thus, the effect of spike, classically captured in an STA as mean signal amplitude in time, may be equivalently described as the *change* of the probability distributions relative to spike, Δ*p*(*x*) [**EQ 1**].

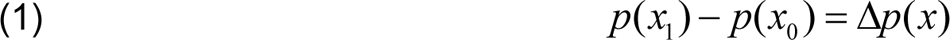

**Figure 4:**
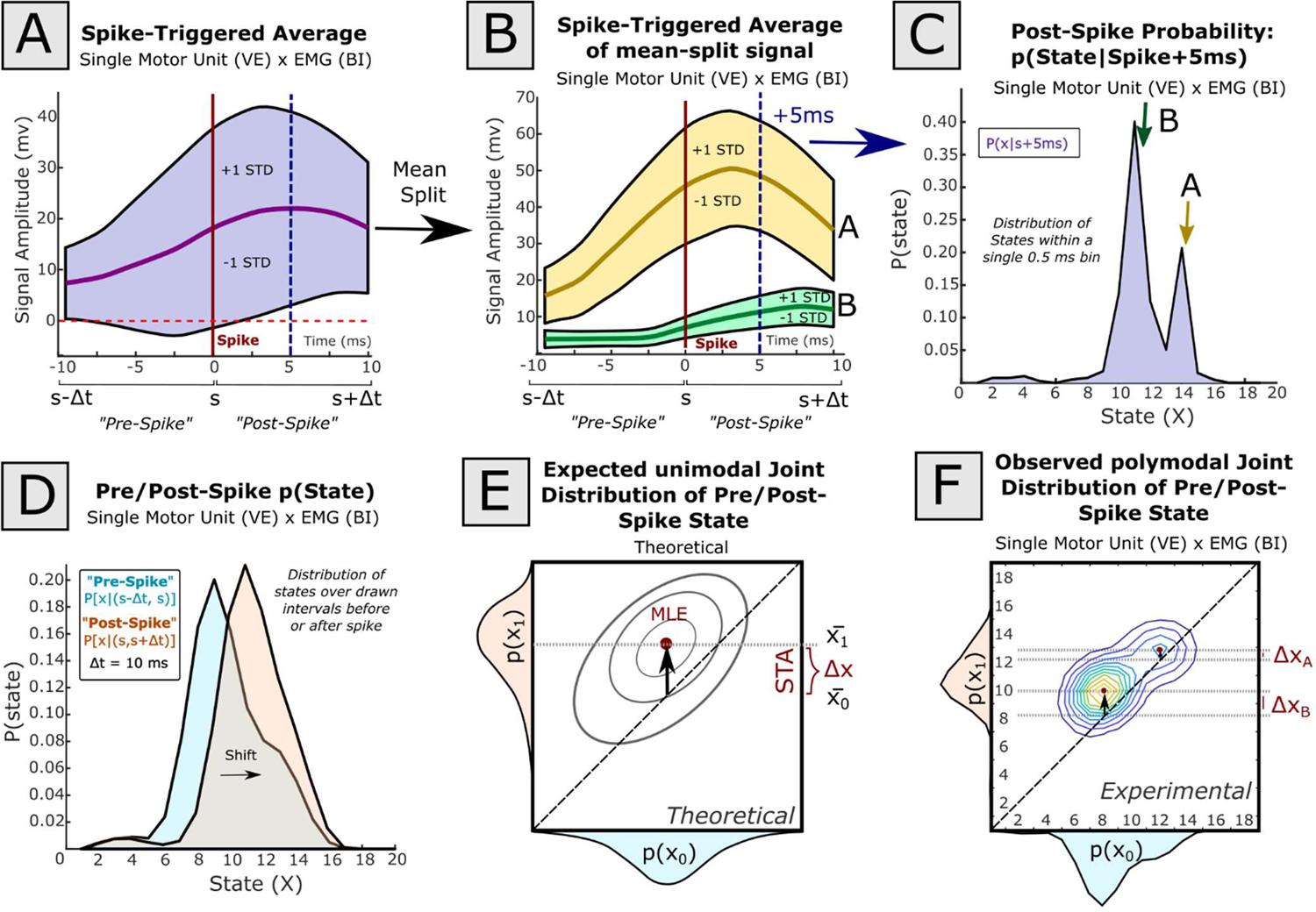
The spike-triggered average (STA) of a signal may vary by background signal behavior. In this example, spiking events from a single motor unit recorded in the vastus externus (VE) muscle is compared against the aggregate EMG signal recorded from the biceps femoris (BI). A window 10 ms before spike to 10 ms after spike was used. **A**) The simple spike-triggered average of BI EMG at spike (red line) within this window has wide variability. The mean STA signal is given as a thick purple line, while ± 1 standard deviation of the windowed signal is described by the breadth of the colored envelope **B)** Separating the aggregate STA by whether the signal is above or below the mean aggregate STA (**A**), provides two STAs (STA-A, STA-B) which exhibit different behavior. **C**) By taking a histogram of binned signal amplitudes (‘states’) at an arbitrary single time point (here, 5 ms after spike; position indicated by dashed blue vertical lines in A and B), this bimodality corresponding to STA-A and STA-B is also seen in state space. **D**) If signal amplitude over the entire perispike interval is quantized into amplitude states, the signal amplitude in the pre-spike and post-spike intervals may be represented as a *pre-spike* and *post-spike* distribution of state. **E**) When *pre-spike* and *post-spike* distributions are described using a joint probability distribution, the theoretical peak value corresponds to the maximum likelihood estimate (MLE) of the post-spike state given pre-spike state. Here, in theoretical data, the STA effect is the migration of the peak from the matrix diagonal (i.e., the average change in state). **F)** If, however, the distribution of *pre-spike* or *post-spike* signal distributions are multi-peaked, as in this physiological example (A-D), transitions to the global peak (X_B_; STA-B) may be suboptimal to describe transitions around local peaks (X_A;_ STA-A). In this example, the best prediction of post-spike state is contingent on whether pre-spike state is closer to 8 or 12.

Practically, because the signal has been quantized into a finite number of states, the *pre-spike, post-spike*, and difference probability distributions are each captured in finite-length vectors. The average *pre-spike, post-spike*, and difference distribution (vectors) could then be averaged across spiking events, as with the STA. In this formulation, within the quantized state space, the maximum likelihood estimation of signal state around spike in state space is given as the global maximum of the joint distribution of the *pre-spike* and *post-spike* state distributions (**Fig. 4E**). Here, the STA corresponds to the movement of the global mean off the joint state diagonal (i.e., the difference in the average state). If, however, *p*(*x*_o_) or *p*(*x*_l_) are not unimodal (as in this example taken from physiological data), local maxima may exist in the joint distribution, independent of the global maximum (**Fig. 4F**). In this case, clearly consideration of the pre-spike state can better inform predictions of *post-spike* state distributions than the average Δ*p*(*x*) alone. <u>In these instances, the STA may perform as a suboptimal</u> <u>description of spike-signal prediction</u>.

#### Stochastic Dynamic Operator Framework

To capture state-dependent relationships in spike-signal correlations more precisely, measures of the changes from *pre-spike* to *post-spike* signal distributions should also be state-sensitive; that is, we desire to know *p*(*x*_l_) = ∑_x_ *p*(*x*_l_|*x*_o_)*p*(*x*_o_), the *post-spike* distribution, given the conditional probability of signal state after spike *and* the prior *pre-spike* signal state distribution.

To map the state-dependent shift in *pre-spike* signal state distribution, we define a linear operator, *L*. *L* is implemented as a matrix which uses the *pre-spike* state distribution *p*(*x*_o_) to generate a column vector providing the *change* in the state distribution, Δ*p*(*x*), <u>given</u> an observed *pre-spike* probability distribution (**EQ2**).

Predictions of *post-spike* distribution conditional on *pre-spike distribution* then may be predicted as *p*(*x*_o_) + Δ*p*(*x*) = *p*(*x*_l_).

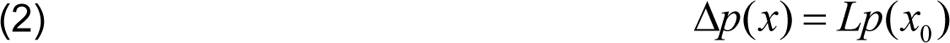

Experimentally, *L* may be recovered as a standard linear estimate across all observed *pre-spike* to *post-spike* distributions. Here, we further elaborate on this process. For a set of *k* spikes, we may define the sampled set of *pre-spike* distributions as *P*(*x*_0_) = [*p*(*x*_0_)_1_…*p*(*x*_0_)*_k_*], the *post-spike* distributions as *P*(*x*_1_) =[*p*(*x*_1_)_1_…*p*(*x*_1_)*_k_*] and the difference distributions as Δ*P*(*x*) = [*p*(Δ*x*)_1_…*p*(Δ*x*)*_k_*]. (For *n* total states and *k* spiking events, this produces n-by-k matrices.) *L* is then estimated as:

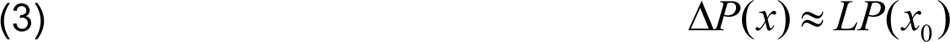

Post-multiplying by the transpose of the *pre-spike* distribution establishes:

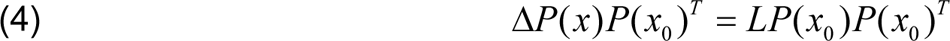

This may be simplified to a standard linear regression problem:

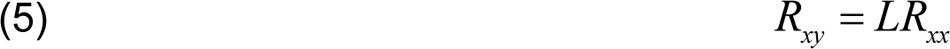

Here *R_xx_* = Δ*P*( *x*)*P*(*x*)*^T^* and *R_0_* = *P*(*x_xy_*)*P*(*x_0_*)*^T^_0_*, and are square matrices of order *n*. *L* is thus derived as:

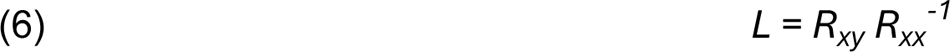

In practice, *L* is a square matrix whose order equals the number of signal states used for analysis. It provides the averaged spike-triggered change in probability of state, given *pre-*spike state, as estimated from the observed data (EQ 3). Thus, *L* is effectively a state-dependent spike-triggered average (albeit for state distributions rather than waveforms). Because *L* operates on probability distributions to describe stochastic signal dynamic changes, *L* is termed a ‘Stochastic Dynamic Operator’ (SDO) (see Sanger, 2011). SDOs were originally developed to describe and manipulate continuous-time stochastic dynamics governed by differential equations. Here, we use the properties of the discrete-time SDO to create a difference equation which describes the state dependence of the spike-triggered average change in probability of state on the prior state.

The matrix operator, *L*, is not uniquely defined. The SDO matrix represents a ‘probability flow’ from the *pre-spike* to the *post-spike* distribution. Probability flows out from each state *i*, to other states j ≠ i. The total outward and inward flow component from each state must sum to 0 to conserve the total probability, and states cannot increase their own probability by flowing into themselves. Four conditions are necessary and sufficient for an SDO matrix to behave as a linear operator (Sanger, 2011): 1) The matrix diagonal must contain non-positive elements, 2) Off-diagonal elements must be non-negative, 3) each column must sum to 0 (if the SDO is modelled as a left-stochastic matrix), and 4) the sum of positive elements within a column must be no greater than 1. From the perspective of a single initial state, *x0=i*, these constraints mean that the *change of probability* of maintaining state at *x1* must be nonpositive (Δ*p*(*x*_l_ = *i*|*x*_o_ = *i*) ≤ 0), while the *change of probability* of transitioning to any other subsequent state must be non-negative; Δ*p*(*x*_l_ ≠ *i*|*x*_o_ = *i*) ≥ 0. (i.e., one may only increase, or sustain, the probability of transitioning to another state and only decrease, or sustain, the probability of maintaining state.)

Having defined, generated, and estimated an SDO across the sampled data, the distribution of states in a *post-spike* distribution given a supplied *pre-spike* prior may be predicted by combination and rearrangement of **EQ1** and **EQ2** (**EQ7**). In this form, the SDO is used to estimate an update to an initial *pre-spike* state distribution.

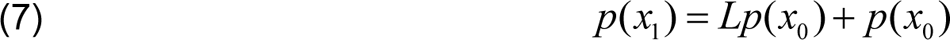

Next, we elaborate SDO analysis as a flexible tool to describe stochastic behavior correlations between physiological data. The SDO methods and analyses described below were generated using the *SDO Analysis Toolkit*, implemented in MATLAB 2023a, running on a Dell Precision Tower 5810 with a Windows 10 operating system.

### Model Hypotheses

To compare the spike-triggered SDO with classical methods, we explored the tested data under several model hypotheses, including the STA. We tested both physiological and model ‘physiological data’ in which ground truth mechanisms originating data were known. Each combination of spiking unit and EMG in our data sets was fit using seven statistical models (below). Each of the 7 hypotheses was embodied as a matrix operator predicting the *change* in the predicted *post-spike* signal state distribution, Δ^-^*p*(*x*), given the *pre-spike* state distribution. (Incidentally, all 7 hypothesis matrices satisfy the constraints to behave as SDOs.) The predicted *post-spike* state distribution was then commonly estimated with the update equation *p*^(*x*_l_) = *p*(*x*_o_) + Δ*p*^(*x*). We then predicted the probability of a post-spike <u>state</u>, or the *post-spike* state <u>distribution</u>, and compared this prediction with the observed values across our datasets. Prediction matrix hypotheses were as follows:

- <u>[H1] Constant</u> Δ*p*^(*x*) = 0: This is the null hypothesis of no effect, and that the data has no statistical drift. The hypothesis matrix is empty (all zeros), hence the predicted *post-spike* state distribution, *p*^(*x*_l_), matches the *pre-spike* distribution, *p*(*x*_o_), exactly. This may occur when signal state changes at relatively low frequency (e.g. no change over the windowed sample), or at random (e.g. white noise), and the spike has no correlation to signal behavior. In terms of the classical STA, this would be equivalent to a flat average waveform in the pre-spike and post-spike interval (i.e., no expected ‘effect’).
- <u>[H2] Diffusing in time</u> Δ*p*^(*x*) = (*G* − *I*)*p*(*x*_o_): (Where, *I* is the identity matrix and G is a Gaussian kernel.) *G,* is convolved with the identity matrix. Consequentially, the predicted *post-spike* distribution reflects diffusion of the *pre-spike* distribution. This is the null hypothesis of no effect but that the data also has a random statistical drift. The predicted *post-spike* distribution has increased variance relative to the *pre-spike* distribution but no change of mean. This may occur if spike is correlated with an unpredictable or randomly fluctuating polymodal spike impulse response, which increases the variance of the *post-spike* distribution when accumulated across time. In both cases, there is no change in the ‘average’ state in the interval. If spike-effects were consistent, there would instead be a clear impulse response in the time domain (e.g., STA).
- <u>[H3] Predicted by spike time alone</u> Δ*p*^(*x*_l_) = (*P*^-^(*x*_l_) − *I*)*p*(*x*_o_): (Where, *I* is the identity matrix.) Here, *P*^-^(*x*_l_) is the average sample *post-spike* distribution. This hypothesis is the classical STA, expressed within state-probability space. Prior state is assumed to have no effect on the resulting spike effect. The predicted change in the *post-spike* distribution causes the *post-spike* distribution to become the average *post-spike* distribution, averaging over all spiking events. (Equivalently, this is the change in the *post-spike* distribution generated from the average *post-spike* waveform.)
- <u>[H4] Predicted only by prior state independent of spike</u> Δ*p*^(*x*_l_) = *L*_B_*p*(*x*_o_): Here, *L*_B_ is an SDO estimated by over *all* time points in the signal, rather than only sampled intervals associated with spike (e.g. the ‘background’ SDO). The same ‘pre-spike’ and ‘post-spike’ interval durations are used for ‘spike-triggered’ responses (although here, all time indices are treated as a ‘spike). This hypothesis matrix permits for signal to have intrinsic (‘background’) dynamics independent of spiking effects and hence demonstrate state-dependent behavior. It can thus be applied to any arbitrary time point of a dynamic signal. This condition may arise, for example, if the system stabilizes a signal around a fixed point, with an effect proportional to deviation (e.g., a spring). In this model the dynamics of state are partly determined by prior state, but spike occurrence has no effect. This also forms the ‘background SDO’ in statistical tests used below, as deviation of the spike-triggered SDO from the ‘background’ SDO indicates a difference in the local system dynamics specifically near the spike occurrence.
- <u>[H5] Predicted by prior state dynamics in the pre-spike interval</u> Δ*p*^(*x*_l_) = (*M*^Δt^ − 1)*p*(*x*_o_): Here, *M*_o_ is a first-order Markov matrix describing state-state transitions between discrete time points only in the *pre-spike* interval for spikes from the sampled neuron. This model is then convolved for the duration of the post-spike interval, Δt, to serve as a transition matrix for prediction. (i.e., generated using the sequence of states in the interval, rather than the distribution of states over the interval. In H5 we estimate local probability dynamics prior to spike as an explanation of post-spike probability dynamics, but independent of the spike event. In this model framework, the sequential state transitions composing the observed time evolutions of state are treated as independent, and independent of spiking.
- <u>[H6] Predicted by prior state, plus a constant effect following a spike</u> Δ*p*^(*x*) = (*L*_B_ + Δ*P*(*x*))*p*(*x*_o_): Here, *L*_B_ is an SDO estimated over *all* time points in the signal (i.e., H4) and Δ*P*(*x*) is the average *change* in the *pre-spike* to *post-spike* distribution, across all samples (i.e. the spike-triggered change). This hypothesis combines the independent state dependent dynamics from H4 with an STA. Here, the STA effect is treated as state-independent after the spike, integrated into the existing, spike-independent, background dynamics.
- <u>[H7] Predicted by prior state and spike (Spike-triggered SDO)</u> Δ*p*^(*x*_l_) = *Lp*(*x*_o_): In this model, the prior state and spike occurrence have interacting effects, and are captured using the spike-triggered SDO matrix, *L* [EQ 2]. In H7, spikes are hypothesized to change the local background signal *dynamics*, which may manifest as different spike-correlated shifts of the *post-spike* distribution and may be state-dependent. For example, a spike may be associated with transitions towards a given state, with directional shifts in the *pre-spike* distribution conditional on whether the signal is ascending <u>or</u> descending towards this state, or with multimodal increased probability of multiple states.

Each model hypothesis examined here assumes a different relationship between a spike source and signal. For example, H4 assumes that the signal state is functionally independent of spiking behavior. H6 assumes that signal state behavior correlates with spike sampling times in a state-dependent fashion, but that spike effects themselves are additive and are not state dependent. H5 assumes that the spike is not a causal agent, although spike occurrence is used to determine epochs where dynamics are sampled. Insofar as the relationship between randomly sampled interneurons and electromyograms may vary, we expected that arbitrary combinations of spike sources and signals in our recorded data could likely be best-modelled by different hypotheses.

### Validation of SDO methods in Simulated Data

We first tested our inference that the spike-triggered average may be insufficient to consistently detect state-dependent or paradoxical interactions within the nervous system using computational models of different stochastic controllers *in silico*. We generated data via well-characterized systems and assessed the specificity and sensitivity of STA and SDO methods to detect these relationships. Eight methods of generating time series data (Y1-Y8), corresponding roughly to the 7 hypotheses (above), and an additional process, were used. To correlate spike-triggered effects, putative spike trains were drawn either randomly or via a Poisson process modulated by a time-varying rate (corresponding to the level of the underlying process).

1. <u>[Y1] Low pass filtered white noise, no control</u>: *Pre-spike* and *post-spike* distributions of state should not systematically vary. No control scheme in use. Spike has no effect (**Fig. 5A**).
2. <u>[Y2] Low pass filtered white noise with spike-triggered random effects</u>: This control is similar to Y1, but at time of spike, there is an added higher amplitude random (white noise) response gated in the 10 ms duration after spiking event. The spike increases signal variance (**Fig. 5B**).
3. <u>[Y3] Low pass filtered white noise with consistent spike-triggered control effects</u>: This is the same as Y1, but with an added consistent impulse response following at time of spike. The spike has a consistent effect (**Fig. 5C**).
4. <u>[Y4] Low pass filtered white noise with stabilizing control dynamics independent of spikes</u>: Here, the signal was synthesized using the update equation *Y*_4_[*t*] = *Y*_4_[*t* − 1] + Δ*Y*_4_[*t*], utilizing the differential of Y1, stabilized towards a fixed point: Δ*Y*_4_[*t*] = *k*(*x*_o_ − *Y*_4_[*t*]) + Δ*Y*_l_[*t*]. Here *X0* is a fixed point (0), and *k* is the stabilizing coefficient (1>*k*>0). Essentially, this equation biases future signal towards zero proportional to the deviation. Spike has no effect (**Fig. 5D**).
5. <u>[Y5] Markov Process independent of spikes</u>: Here, a sequence was generated via a random walk of a Markov matrix. The signal was mean-leveled around zero. Spike has no effect (**Fig. 5E**).
6. <u>[Y6] Stabilizing Dynamics with consistent spike-triggered effects</u>: This is the same as Y4, with the spike control impulse responses of Y3 added. Spike has a consistent effect (**Fig. 5F**).
7. <u>[Y7] Stabilizing Dynamics with spike-triggered dynamics</u>: This is the same as Y4, however, in the 10 ms duration after the spike, a second stabilizing equation is additionally activated (of the same form in Y4). For the spike-triggered dynamics, the equilibrium of the control is positive, *X0* > 0, such that background dynamics and spike-triggered dynamics stabilize towards different points. A spike activates a consistent change in dynamics (**Fig. 5G**).
8. [Y8] <u>Autoregressive Integrated Moving Average [ARIMA] Stationary Stochastic Process</u>: ARIMA models are commonly used to parsimoniously describe and forecast stochastic time series data. Here, a stationary ARIMA(2,2) model was used. The signal was mean-leveled around zero. Spike has no effect (**Fig. 5H**).

**Figure 5:**
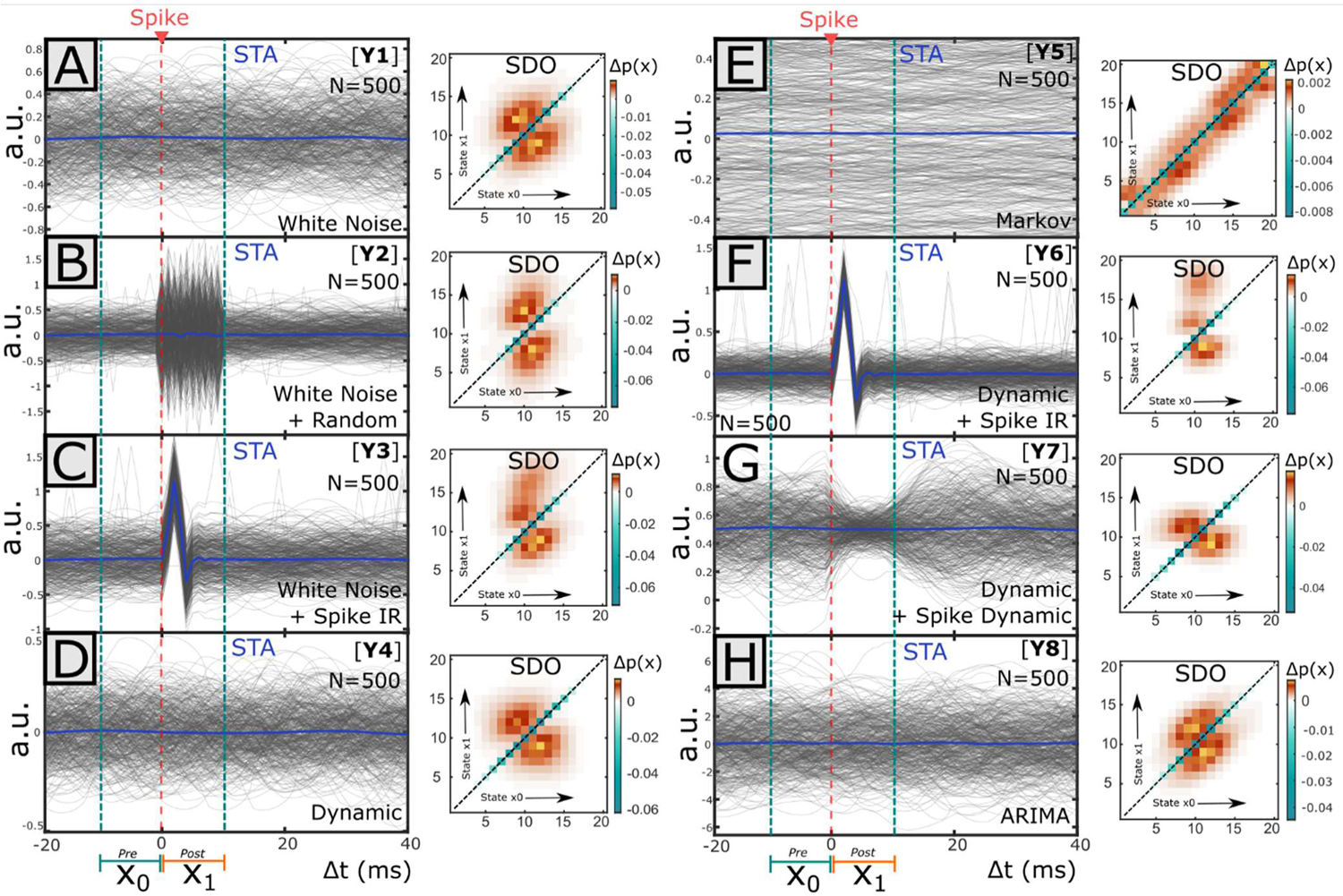
Comparison of the STA impulse response and SDO matrices obtained from simulated stochastic controls and data Time series and point-process data was simulated with different stochastic processes and spike-signal control relationship (or lack thereof). For each simulated spiking event, a sample of time series data was extracted in the window 10ms before spike to 10 ms after spike. The STA impulse response was estimated as the average signal across all spike-triggered samples. The pre-spike window was defined as 10ms before spike until the time of spike, and the post-spike interval as the 10 ms following the spiking event. In each case, the SDO was generated using the distribution of states extracted across each interval. 20 states were linearly defined from the minimum and maximum of the observed signal values. Time series were stationary about a mean value (usually zero), with similar amplitude ranges on either side of the mean. Hence, the average signal level corresponds to an intermediate signal state (i.e., 10-11), and SDOs had values around the middle of the matrix. Representative examples of STAs and SDOs from simulated data are plotted here. The 500 spike-triggered window signals are overlaid as gray traces in time. **A**) [Y1] Time series was generated using lowpass filtered white noise. Spikes have no effect. STA is flat and SDO is symmetrical about the diagonal. **B**) [Y2] Spikes evoke increased noise level impulses in otherwise white noise. STA is flat, while the SDO is elongated in the horizontal dimension (corresponding to the increased signal variance after spike). **C**) [Y3] Spikes have consistent effects within a white noise signal. STA extracts the impulse response. The SDO is elongated above the matrix diagonal, showing an increase in transition towards higher states (amplitudes). **D**) Signal is generated by a stabilizing dynamic equation. Spike has no effect and STA is flat. The SDO demonstrates more stabilizing effects than in white noise (Y1); above-average input states (11-15) are pushed towards lower states (5-10) while below-average input states (5-10) are pushed towards higher states (10-15), as indicated by SDO elements. **E**) [Y5] Markov process. Subsequent signal amplitude transitions to similar amplitude as prior amplitude. Spike has no effect. STA is flat. The SDO indicates that changes in signal distribution at spike time are minor and bidirectional (indicated by the limited band on the matrix diagonals). **F**) [Y6] Spike has consistent effect within a process described by a stabilizing dynamical equation. As with Y3, STA extracts the impulse response and SDO is elongated. **G**) [Y7] Spike induces dynamic effects, while signal is also generated by a stabilizing dynamic equation. STA is flat. SDO is elongated in the horizontal dimension, indicating pre-spike signal variance is less than post-spike variance. **H**) [Y8] Signal is generated from an ARIMA(2,2) process. STA is flat. As with the white noise (Y1), the SDO is relatively symmetrical about the matrix diagonal.

We compared the STA and SDO methods of analyzing data from these various simulations of model systems in which spikes were independent of dynamics or participated in the control of dynamics.

### Computational Implementation of STA Methods

We used three methods of calculating the spike-triggered average between spike and (simulated) rectified EMG, and one measure of spike tuning. A spike-triggered effect was considered ‘significant’ if any test of effect returned a positive result using α = 0.05. To avoid spurious correlations, only spike trains with a minimum of 500 spikes were included for analysis.

#### Simple Effects

This is a simple test of correlation in signal amplitude relative to spike correlations. The average signal amplitude in each 20ms pre-spike interval and the 20ms post-spike interval was captured for all spiking events. We then tested the null hypothesis that the difference in the distribution of average signal amplitude in the pre-spike and post-spike interval was normally distributed around zero (i.e., a paired t-test).

#### Detrended Effects

The increment-shifted average (ISA) technique attempts to isolate spike-correlated effects in signal from potentially slower curvilinear trends in signal amplitude preceding and following spike (Davidson, et al, 2007). In this framework, for every spike, *s*, a segment −40ms prior to and 100ms after the spike is extracted from the signal envelope. For every discrete time increment *t* ∈ [−20,40] around spike, a sample waveform x[s+t] is collected, and the discrete time increment t is shifted right (i.e., increased by 1). The average of these waveforms is the increment-shifted average (ISA). Subtraction of the increment-shifted average waveform from the spike-triggered signal in the window 20ms prior to and 40ms after spike may better isolate spike-correlated events relative to slower-resolving background behavior. The mean of the pre-spike and post-spike expected baseline windows (20-0 ms before spike; 20-40 ms after spike) is then subtracted from the mean of the ISA-subtracted window in the test interval (0-20 ms after spike), isolating the average change of signal specific to this interval (Poliakov and Schieber, 1998). We then tested the null hypothesis that the resultant distribution of spike-effects is normal and centered around zero.

#### Bootstrapped Deviation

The Chronux MATLAB package (Mitra & Bokil, 2008; Bokil, et al., 2010) similarly tests for significant spike-triggered average relationships by collecting pre-spike (20-0ms before spike) and post-spike (0-20ms after spike) windows. The standard deviation of relative signal amplitude across the population of *N* spikes at every relative time index, *t*, is estimated using the average waveforms from 20 bootstrapped populations of *N* spike indices (i.e., resampling with replacement). A spike-triggered average relationship was considered significant if the rectified, mean-leveled, STA waveform had 1 or more points above threshold anywhere within the interval T = [-20ms, +20ms] relative to spike. Threshold was defined by calculating the inverse t-statistic after Bonferroni correction to the familywise error rate for the number of points tested. (This corresponded to ∼2.99 standard deviations for 40 points).

#### Computational Implementation of SDO Methods

To generate, visualize, and analyze SDOs, we developed the *SDO Analysis Toolkit*, written in the MATLAB programming language. For each time series signal (Y1-Y8 in simulated data, or rectified aggregate EMG in the physiological data), signal states were defined as 20 intervals on between the channel-wise minimum and maximum values of signal amplitude observed over the entire data set. (Note that simulated data used a linear scale while the EMG data used a logarithmic scale.) Under the SDO model, each spike was treated as an independent event. For each spike, we extracted signal states in a short interval (10 ms) before (“pre-spike”) or after (“post-spike”) the spike event. The frequency histogram of states within these observed intervals relative to spike comprised the *pre-spike* and *post-spike* probability distributions. The *pre-spike* distribution included the time bin containing the timestamp as the final element.

#### Identifying Significant SDOs

Spike-triggered SDOs were tested for significant spike-signal correlations using Monte Carlo sampling. We describe this process as follows: First, we generated 1000 shuffled spike trains, for each observed spiking unit (e.g., neuron). We included two methods of shuffling spike times, both included within the *SDO Analysis Toolkit*. The first is to directly shuffle the observed interspike intervals on a trial-wise basis, thus preserving the durations between impulses. The second method is to estimate the rate process underlying the observed spike train, assuming spikes were generated by a renewal process, then to resample this process 1000 times. (The *SDO Analysis Toolkit* allows for various filtering kernels for this purpose.) In both cases, the 1000 ‘shuffled’ spike trains from the tested spiking unit were then used to produce 1000 ‘shuffled’ spike train SDOs, using the same signal as the tested experimental SDO.

In both spike-triggered and shuffled-spike SDOs, we measured four features of SDO effect, and one feature of state-tuning:

1. Matrix element values: The sum of squares error (SSE) calculated between each element of the spike-triggered SDO matrix vs. the distribution of values for that element in the shuffled distribution.
2. Matrix similarity: Overall cumulative SSE of the spike-triggered SDO matrix and the shuffled SDO matrices.
3. State-wise *post-spike* distribution directional shift/bias: For each input state (SDO column), the sum of the elements above the matrix diagonal vs the elements below the matrix diagonal. (This corresponds to the expected change in mean imposed by the SDO, conditional on input state).
4. Array-wise total *post-spike* distribution directional shift/bias: The sum of all SDO matrix elements above the diagonal vs. the elements below the diagonal (i.e., does the spike coarsely facilitate or inhibit signal?)
5. Pre-spike state tuning: The Kullback-Leibler Divergence between the observed state at spike, *p*(*x*|*s*), vs. shuffled spike times. (Note that this is not a measure of spike effect, but can be used to identify significant relationships between signal level and spike occurrence.)

The significance of the spike-triggered SDO was determined by comparing the spike-triggered test statistic vs. the null distribution of test statistics from the 1000 shuffled data-derived SDOs. Here, the difference between the spike-triggered SDO statistic and the mean of the shuffled-spike SDO statistics was compared with the internal variance of the population of 1000 ‘shuffled’ SDOs (i.e., is the test result from the spike-triggered SDO significantly different than would be predicted from the shuffled distribution?) A P-value of < 0.05, with a Bonferroni correction for the number of states (20), was used to determine significance.

#### Matrix Normalization by Prior Distribution of State

The SDO matrix describes the average change of state, given state. When averaged across all spiking events, SDO estimation is influenced by the underlying joint distribution of pre-spike and post-spike state describing the overall frequency of state observations (**Fig. 6A**). Thus, the columns of the default SDO are scaled proportional to the prior (i.e., *pre-spike*) distribution of states (**Fig. 6B**). To account for differences in the sampling of pre-spike state (column scaling) between the background, ‘shuffled’, and spike-triggered SDOs, the columns of the SDO matrices were first normalized, then rescaled before evaluating significant differences. To normalize an SDO matrix, each column *j* of the SDO is scaled by the reciprocal of the probability of state *j* in the *pre-spike* distribution, 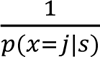. (When applied to the joint distribution of state *p*(*x* |*x*), this is equivalent to normalizing each column to 1 (**Fig. 6C**).) In this normalized form, the SDO represents the conditional *change* of the probability of state, *given* an initial state distribution, Δ*p*(*x*_o_|*x*_o_), and hence emphasizes magnitude of correlated signal change, by state (**Fig. 6D**). The normalized (conditional) SDO matrix is the form used for generating predictions of *Δp(x)* as in EQ2.

**Figure 6:**
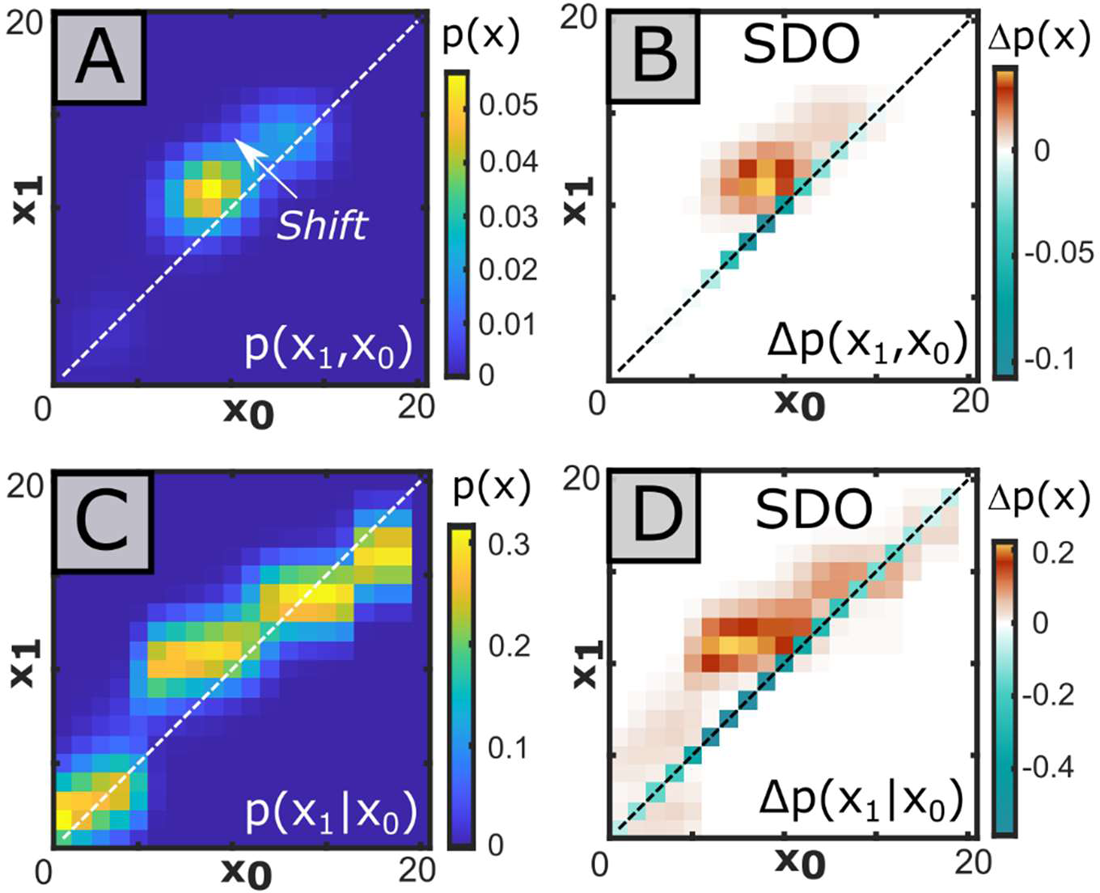
Stochastic Dynamic Operators alter background state transition matrices to compose spike-triggered transition matrices. In this example, *pre-spike* and *post-spike* distributions of signal state were generated from the spikes of a single motor unit in the vastus externus against the analog EMG signal recorded in the biceps femoris, using a time interval of 10 ms for both the *pre-spike* and *post-spike* distributions. **A)** The spike-triggered average joint distribution of the pre-spike and post-spike state distributions behaves as the description of state transitions around spike. **B**) A spike-triggered SDO captures the *change* of state probability across the 10ms *pre-spike* and *post-spike* distribution of states, given an occurrence of spike. This SDO shows strong directional effects, evidenced by the asymmetrical banding relative to the diagonal. **C)** Normalization of each column of the joint distribution to 1 creates a left transition matrix, representing the expected probability of *post-spike* state distribution given a *pre-spike* state distribution. **D**) Normalizing the columns of the SDO by the same factor used in the joint distribution [C], creates the normalized SDO. This form is used to predict the conditional change of state distributions.

Column-wise rescaling of the normalized SDOs by a new expected *pre-spike* distribution of state reparametrizes the matrix with a new prior distribution. In the (re)parameterized form, the SDO emphasizes the correlated change in probability of signal state by expected frequency of observation of initial state (i.e., prior distribution of state). The parameterized form of the SDO is preferred for plotting and evaluating significance, as this emphasizes the expected contribution of the SDO, weighted by the probability of observing the change. The ‘spike-shuffled’ and background SDOs were reparametrized to the *pre-spike* distribution of the spike-triggered SDO prior to performing statistical tests of significance as mentioned above. (This mitigates the occurrence of ‘significant’ differences spuriously occurring from differences in state sampling.) Both the normalized and parameterized SDOs can be readily plotted within the *SDO Analysis Toolkit*.

### Visualization of SDO and SDO Effect Predictions

Visualization of the correlation between spikes and signal is important for intuition. The spike impulse response of the STA may be readily visualized and interpreted from the average time-varying waveform of sampled signal amplitude across all spiking events. In contrast, the SDO describes the change in state probability distributions without prescribing a specific temporal waveform structure. Thus, any time effects would not be immediately apparent from the *pre-spike* and *post-spike* distributions for the SDO. Nonetheless, considering the underlying state transition probability and signal behavior over time is important.

We observed that the spike-triggered response may vary by signal state at time of spike (**Fig. 7A**). To identify coarse signal behavior relative to spiking events in the SDO framework, we developed the spike-triggered impulse response probability distribution (STIRPD) description (**Fig. 7B**). The STIRPD is effectively a transformation of the classical STA into the probability state space but has the advantage of more robustly describing the relationship between the spiking source and signal behavior in time. To calculate the STIRPD, as with the STA, for each spiking event, *s,* the state-quantized signal is sampled on an interval around spike from (*s* − Δ*t, s* + Δ*t*). Subsequently, the distribution of signal states at every time point *t, p(x,t)*, is calculated from the same relative time bin across all sampled signals. The value of each element represents the probability of observing a given state (as opposed to another state) at a given time, *STIRPD*_ij_ = *p*(*x* = *i*|*t* = *j*).

**Figure 7:**
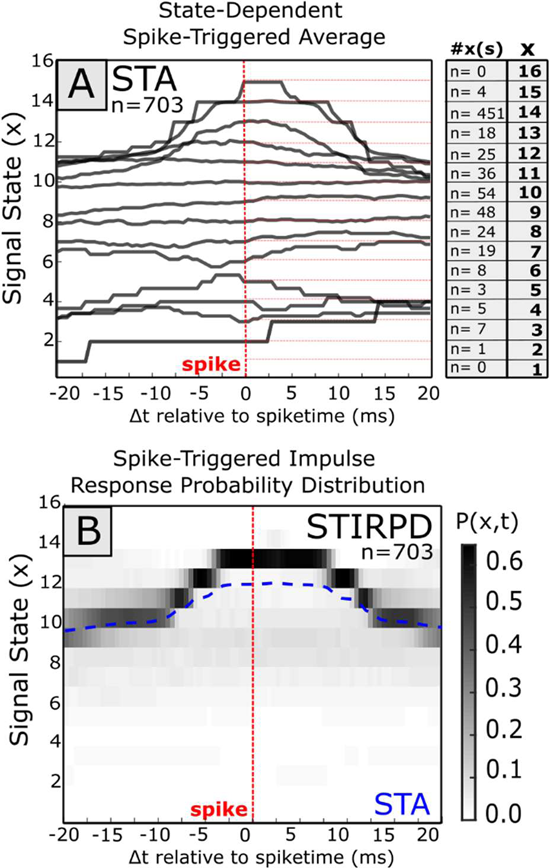
Expressing STAs and probability distributions. The spike-triggered impulse response probability distribution (STIRPD) is a combination of the classical spike-triggered average and state probability distributions. **A**) The relationship between the spike train of an interneuron is compared against EMG (filtered acausally). In the state-dependent STA, a set interval [s-Δt, s+Δt] of signal behavior is extracted for every spiking event (s), sampled in discrete time. A state, *x,* is assigned to every observed time point, t, (*t* ⋲ [*s* − Δ*t, s*Δ*t*]) within this interval to provide a sequence of states, *x*(*t*). When segregating the STA by state at time of spike, *x*(*s*), state dependent behavior may be observed. **B)** The spike-triggered impulse response probability distribution extends the STA by calculating the distribution of states at each measured time point *p*(*x*, *t*) across all spiking events. Because both states and time are discrete, the STIRPD is a finite matrix and may be displayed as an image. Here, the ‘arching’ signal behavior present at the higher states in the STA is present as a dark band, but the STIRPD nonetheless captures the relatively invariant signal behavior in states 6-10 in the perispike interval. **C**) Each element of the STIRPD is the observed probability of a particular signal state at a particular time relative to spike. Each column of the STIRPD contains the full probability mass function at the time point, while the STA alone would only correspond to the mean of this distribution. The STIRPD is thus a more complete representation of the STA, which may be useful for identifying gross signal behavior in the perispike interval. **D**) Summing over the STIRPD over the pre-spike and post-spike time columns provides the matching *pre-spike* and *post-spike* state distributions used to generate the joint probability state matrix and the stochastic dynamic operator. Horizontal separation of state mass in the STIRPD (i.e., banding) will result in multimodal pre-spike or post-spike distributions elements of the joint probability matrix and may be indicative of cases where the STA is insufficient. Here, the STA is overlaid on the STIRPD in blue.

Because both time and state are defined discretely, the STIRPD may be displayed as a raster image and interpreted qualitatively. The familiar spike-triggered average (in state space: mean state) is laid over the probability distribution. The probability distribution of state at time of spike, *p(x|s)*, is captured as a single column vector for the time bin containing the spike on the STIRPD and thus can reveal if the distribution is expected to be unimodal or multimodal, or if there is a coarse signal behavior change around spike. The summation of state probability over time in the STIRPD during the pre-spike interval, [*s* − Δ*t, s*], provides *p*(*x*_o_), while summation of the STIRPD over the post-spike interval, provides *p*(*x*_l_), used in the calculation of the SDO. This description thus links the SDO and STA visualization methods.

As with the STIRPD, we could also qualitatively infer the relationship between a target signal and spike source neuron for the pre-state (x0) to the post-spike (x1) state using visualizations of the SDO matrix directly (**Fig. 8A**). When inspecting the SDO, we also found it intuitively helpful to plot a shear of the SDO matrix such that the prior matrix diagonal, corresponding to (*i* = *j*) is aligned horizontally (**Fig. 8B**). This transformation emphasizes the direction of the SDO effects, Δ*p*(Δ*x*, *t*): Elements above the shear SDO horizontal correspond to transitions towards higher states (*x*_l_ > *x*_o_), while below-diagonal corresponds to transition towards lower states (*x*_l_ < *x*_o_). The sign and magnitude of these elements may be used to interpret SDO effect for a given input state. (e.g., for an input state which has positive elements (increased probability) above the shear-horizontal “towards higher states” and negative elements (decreased probability) below the shear-horizontal “towards lower states”, the net effect for that input state is a shift towards the higher states.)

**Figure 8:**
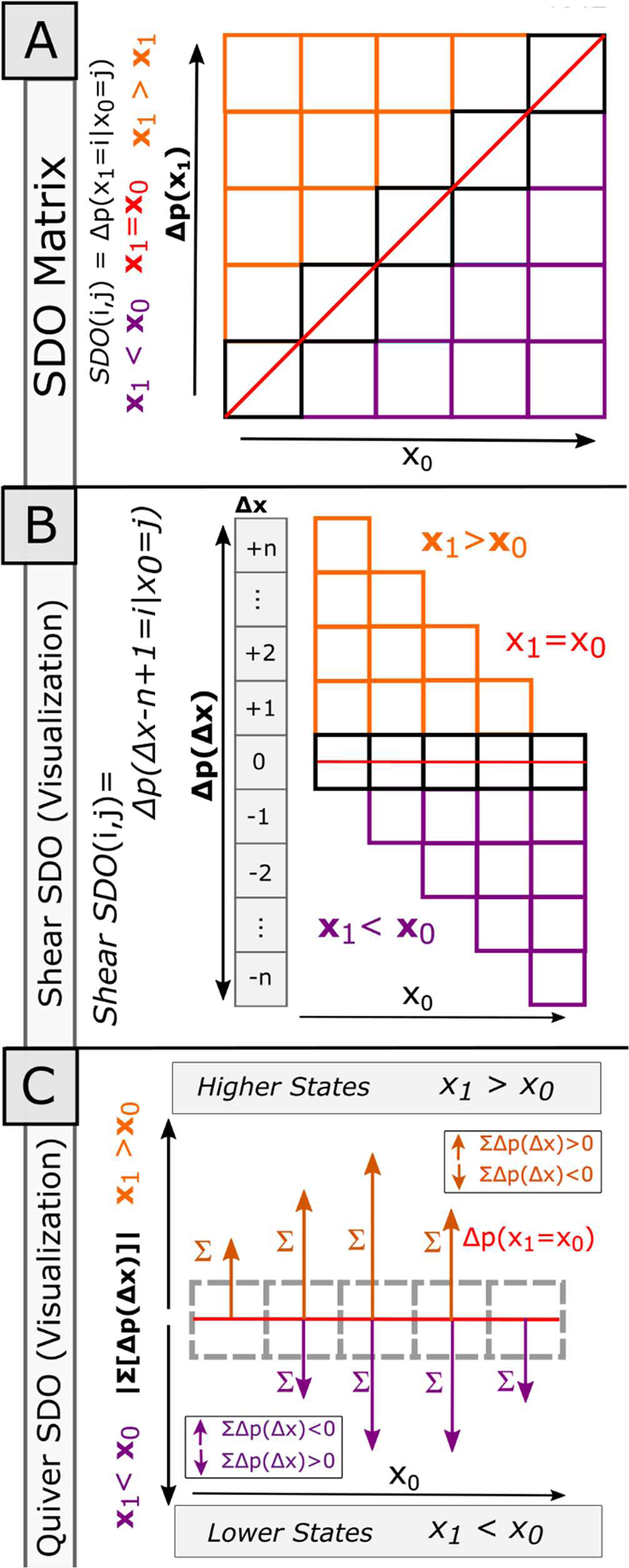
Alternative Visualizations of the SDO Matrix. The effect of a spike-triggered SDO on the probability of signal state between two observations *x_0_* and *x_1_* may be qualitatively inferred from the SDO directly. **A)** The SDO matrix may be divided into three regions, the primary diagonal (red line), representing the probability of maintaining state; the ‘above-diagonal’ (orange) region, representing the probability of increasing state; and the ‘below-diagonal’ (purple) region, representing the probability of decreasing state. **B**) The square SDO matrix may be sheared to orient the prior matrix diagonal to the horizontal. The shear SDO visualization emphasizes the change in probability of state, Δp(x), relative to the change of state, Δx. As SDO operators are sparse matrices and usually concentrated around the diagonal when time intervals are short, the shear SDO may be trimmed to a few rows to compactly display the effects of an SDO. **C**) The magnitude and directional effects of an SDO can be coarsely described by summing Δp(x_1_|x_0_) separately for the regions above (orange) and below (purple) the SDO matrix diagonal. In the quiver SDO visualization, these column sums are represented as vertical vectors oriented above or below the abscissa, respectively. The length of each vector for state *x* is the magnitude of the summed change in probability. If the sum is positive, the vector points away from the abscissa, and towards the abscissa if it is negative. The elements of the diagonal are plotted as a line within the same axes. The orientation and magnitude of vectors for each state, x, provide a coarse measure of influence of Δp(x_1_|x_0_).

A third visualization of the SDO we explored is the ‘Quiver’ representation of an SDO (**Fig. 8C**), which emphasizes the magnitude and coarse directional biases of Δ*p*(*x*) for a given input state. Here, in the quiver representation, each column of the shear SDO matrix is summed over (1) the above-horizontal elements and (2) the below-horizontal elements, and each represented by a vector whose magnitude is proportional to this sum, originating at the abscissa, and projecting up and down, respectively. The elements of the reference horizontal row (the original SDO matrix diagonal), corresponding to Δ*p*(*x*_l_ = *x*_o_), are co-plotted by a simple line (because the SDO matrix diagonal is defined to be non-positive, this trace is negative). These vectors indicate the coarse directional effect of the SDO for a given pre-spike state: If both vectors above and below have an equivalent magnitude for a state, then the change in post-spike state overall is not directional (i.e., there is no change in the mean of the post-spike state distribution contributed by this input state), while unbalanced vectors indicate a coarse direction of change in the post-spike predicted state distribution.

### Assessing the Prediction Utility of Significant SDOs

Finally, we tested the capability of the SDO to predict post-spike signal behavior within our datasets. We assessed two types of prediction: **1**) Predictions of the *post-spike* state *distributions*, *p*^(*x*_l_), and **2**) predictions of most likely *single* state within the same post-spike interval, *x*-_l_ = *argmax*_xi_: *p*^(*x*_l_ = *x*_i_). Predictions of the *post-spike* state distribution (1) capture overall signal dynamics, while predictions of single post-spike states (2) reflect utilization of such a prediction towards directing an exclusive action or decision (e.g., ‘go’/’no-go’, tracking-target, etc.). (Note that even in instances where the spike-triggered SDOs did not significantly differ from shuffled-spike or background, SDOs could still be used to predict signal behavior, but we have limited our scope here to significant cases to compare the STA vs. the SDO directly.)

When evaluating predictions of an SDO (that is, the correlation between a particular spiking unit and signal), the observed *pre-spike* and *post-spike* distributions of signal state were extracted for each spiking event. The predicted *post-spike* state distribution, *p*^(*x*_l_|*H*), were generated on an event-wise basis for each of the 7 matrix model hypotheses [H1-H7] described above, (including the spike-driven SDO as H7), in accordance with EQ 7. The ‘single best state’ of each predicted *post-spike* distribution was defined as the most-likely state (i.e. peak). When evaluating model-predicted *post-spike* distributions, the experimentally observed *post-spike* distribution, *p*(*x*_l_), served as ground truth. Hypotheses which minimized the reconstruction error of the *post-sspike* distributions with the ground truth model were considered the model of best fit (*argmin*_H_: *p*(*x*_l_) − *p*^(*x*_l_|*H*) for H1-H7). (Note that here, the H3 matrix is used to predict the effects of the classical STA represented in the probability space, corresponding to the average change in signal distributions which would otherwise be predicted from the time-varying envelope.) Spiking events, and associated errors, were treated as independent observations. The difference between predicted and observed *distributions* was quantified using the Kullback-Leibler Divergence (KLD) and log-likelihood. The difference between predicted and observed *single states* was quantified using the error frequency (e0), absolute error (e1), and squared (e2) error. Because the spike-wise error rate for these predictions to single states are integer values, and the population of errors is not expected to resemble a continuous distribution (and hence may be ill-suited for tests of means or medians), we instead calculate the cumulative error and bootstrap this statistic to evaluate the confidence intervals. If the distribution of two model hypothesis errors did not overlap at the 95% confidence interval, they were considered significantly different. Observed error rates were considered significantly different if there was no overlap of the estimated 95% confidence intervals, corresponding to a p-value of < 0.05. We used Cohen’s D statistic (Cohen, 1988) to measure the effect size between bootstrapped cumulative error distributions. Additional information about these statistical methods may be found in the Appendix. We hypothesized and expected that, by capturing information of the spike effect more effectively, the spike-triggered SDO (H7) predictions would be the best most of the time.

## RESULTS

### SDO Methods are more Sensitive to State-Dependent relationships than the STA

Within the simulated data, the white noise (Y1), background dynamics (Y4), Markov (Y5), and ARIMA (Y8) stochastic processes harbored no spike-triggered effects. We considered the significance of spike-triggered tests against these processes as false positives. Similarly, when spiking events induced a change in signal amplitude or dynamics (Y3,Y4,Y6,Y7), we considered the failure to detect significance as false negatives. Significance tests of the STA and SDO had different performance by condition (**Table 1**). The t-test method reported Y2 (random impulses) as significant 100% of the time, while both the ISA-subtraction and Chronux-based STA methods never reported Y2 as significant (The desirability of this outcome may vary by circumstance). Only SDO measures could reliability detect significance of Y7.

**Table 1:** Performance of the different significance tests of spike-triggered effects for the STA and SDO methods over 1600 simulations. Test performance as evaluated as the proportion of ‘correct’ tests to the total number of tests, by method and condition. Here, Y1 (white noise), Y4 (background dynamics), Y5 (Markov process), and Y8 (ARIMA) were not influenced by spike, hence failure to reject the null hypothesis (i.e., no spike effect) was treated as ‘correct’. For Y2 (spike-triggered random impulses), Y3 (spike-triggered consistent impulses), Y6 (spike-triggered consistent impulses on a dynamic background) and Y7 (spike-triggered dynamics), a significant spike-triggered effect was treated as ‘correct’. The average performance is the average value across each row. Here, SDO methods are highly sensitive and specific across all tested data modalities.

| Basis | Test | Y1 | Y2 | Y3 | Y4 | Y5 | Y6 | Y7 | Y8 | Average Performance |
| --- | --- | --- | --- | --- | --- | --- | --- | --- | --- | --- |
| STA | Simple t-test | 0.95 | 1.00 | 1.00 | 0.95 | 0.94 | 1.00 | 0.09 | 0.96 | 0.86 |
| STA | ISA-Subtracted | 0.75 | 0.00 | 1.00 | 0.81 | 0.88 | 1.00 | 0.33 | 0.83 | 0.70 |
| STA | Chronux | 1.00 | 0.00 | 1.00 | 0.99 | 1.00 | 1.00 | 0.07 | 1.00 | 0.76 |
| SDO | Matrix Elements | 1.00 | 0.90 | 0.86 | 1.00 | 1.00 | 0.89 | 0.38 | 1.00 | 0.88 |
| SDO | SDO Similarity | 0.99 | 0.90 | 1.00 | 0.98 | 1.00 | 1.00 | 0.90 | 1.00 | 0.97 |
| SDO | P(x0,x1) Similarity | 0.99 | 0.90 | 0.91 | 0.89 | 0.90 | 1.00 | 1.00 | 0.90 | 0.94 |
| SDO | Total Coarse Bias | 0.97 | 0.80 | 0.88 | 0.97 | 1.00 | 0.90 | 0.89 | 0.99 | 0.92 |

We then assessed the relative *sensitivity* (i.e., the probability that a ‘True’ effect is classified as ‘True’) and *specificity* (i.e., the probability that a ‘False’ effect is classified as ‘False’) of the STA and SDO methods in simulated data. We assessed STA and SDO significance across all measures: Here, a STA or SDO was considered “significant” if any test of significance of spike effect returned significance (as in an initial assay for significant relationships in experimental data). When describing simple relationships between spike and signal (e.g., Y3; noise convolved with consistent impulse responses), STA and SDO methods were equally capable of identifying consistent spike-triggered responses (Y3, Y6) throughout simulated data. This sensitivity was upheld even when using random impulses (Y4). The STA had a higher false positive rate for time series data generated from a Markov process (Y5) than the SDO. However, in the case where spiking events caused a change in local system dynamics (Y7), the STA was significantly less sensitive than the SDO, 39.3±4.9% [STA]^a^ vs. 99.8±0.4% [SDO]^b^ (1-tailed t-test, p<0.00001).

We also observed a difference in specificity with the STA and SDO significance measures depending on the origin of stochastic signals (Y1-Y8). When evaluating the grand mean across all conditions, STA had a significantly higher number of false-positive significance tests in simulation for white noise (Y1), 27.7±3.1% [STA]^c^ vs. 15.4±6.8% [SDO]^d^ (1-tailed t-test, p<0.00001), signals with background dynamics (Y4), 22.3±3.3% [STA]^e^ vs. 17.9±8.4% [SDO]^f^ (1-tailed t-test, p=0.029), and Markov processes (Y5), 17.1±5.6% [STA]^g^ vs 2.7±3.4% [SDO]^h^ (1-tailed t-test, p<0.00001). The specificity between the STA and SDO methods for the ARMA(3,2) process (Y8) did not significantly differ 19.1±4.4% [STA]^i^ vs. 15.4±9.8% [SDO]^j^ (1-tailed t-test, p=0.12).

We first tested the importance of the number of spikes vs. the number of bootstrapped ‘shuffles’ used for SDO significance estimation. (Note that in this case, the number of shuffles, used for evaluating SDO significance, is expected to have no effect on the STA. For each simulation, 8 stochastic signals [Y1-Y8] and a common spike train were generated, as described above. 100 simulations were performed for every combination of number of spikes (250, 500, 1000, 2000), and number of shuffles (250, 500, 1000, 2000). For these simulations, other variables (e.g., filtering, number of states, length of pre-spike/post-spike intervals, signal transformations, amplitude/shape of the spike-triggered impulse response) were kept consistent. The battery of SDO significance measures remained highly sensitive with all tested parameters (**Table 2**) and, unsurprisingly, became more specific as additional bootstraps were used for estimating the distribution of the null statistic for SDO (**Table 3**). As expected, measures of STA specificity did not vary by number of shuffles used for SDO significance estimation (**Table 4**) For all combinations, except 250 spikes with 250 bootstraps, SDO methods outperformed STA measures of significance (**Table 5**). Having observed that the SDO could provide additional sensitivity and specificity compared to the STA in simulated data we set about applying the SDO to physiological data.

**Table 2:** SDO Significance Test Sensitivity by Number of Shuffles and Spikes: Proportion of 100 simulations (per condition) where *any* of the four SDO significant tests reported a significant spike-signal relationship, out of all simulations with a spike-signal relationship (Y2, Y3, Y6,Y7), by number of spikes and bootstraps. Here, the number of spikes and bootstraps did not affect the sensitivity of the SDO significance measurements across the simulated data.

| SDO | #Spikes | 250 | 500 | 1000 | 2000 |
| --- | --- | --- | --- | --- | --- |
| #Shuffles |  |  |  |  |  |
| 2000 |  | 1 | 1 | 1 | 1 |
| 1000 |  | 0.99 | 0.99 | 1 | 1 |
| 500 |  | 1 | 1 | 1 | 1 |
| 250 |  | 1 | 1 | 1 | 1 |

**Table 3:**
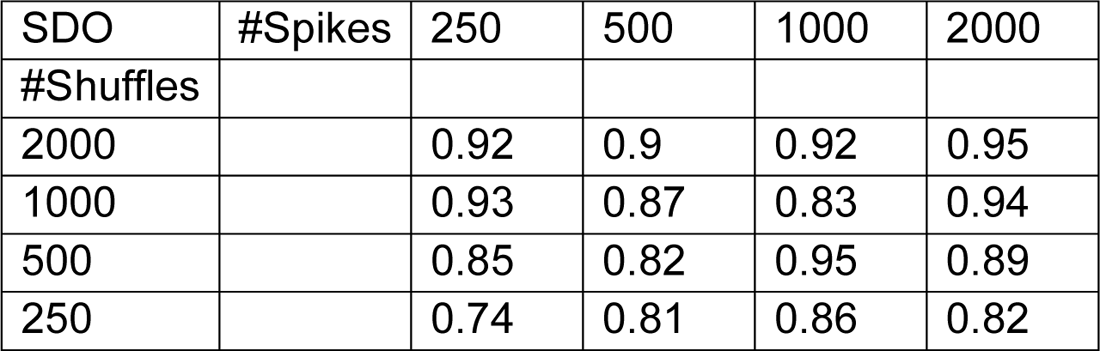
SDO Significance Test Specificity by Number of Shuffles and Spike: Proportion of 100 simulations (per condition) where *none* of the four SDO significant tests reported a significant spike-signal relationship, out of all simulations where spike had no effect (Y1, Y4, Y5, Y8). Here, unsurprisingly, the specificity of the SDO methods increase with a higher number of shuffles used for generating the null distribution for significance tests.

| SDO | #Spikes | 250 | 500 | 1000 | 2000 |
| --- | --- | --- | --- | --- | --- |
| #Shuffles |  |  |  |  |  |
| 2000 |  | 0.92 | 0.9 | 0.92 | 0.95 |
| 1000 |  | 0.93 | 0.87 | 0.83 | 0.94 |
| 500 |  | 0.85 | 0.82 | 0.95 | 0.89 |
| 250 |  | 0.74 | 0.81 | 0.86 | 0.82 |

**Table 4:** STA Significance Sensitivity: Proportion of 100 simulations (per condition) where *any* of the three STA significant tests reported a significant spike-signal relationship, out of all simulations with a spike-signal relationship (Y2, Y3, Y6,Y7). Here, the only the number of spikes vary in the STA significance calculations (shuffles counts are only used for co-estimated SDO significance). In this instance, the sensitivity of STA significance methods does not vary by spike count.

| STA | #Spikes | 250 | 500 | 1000 | 2000 |
| --- | --- | --- | --- | --- | --- |
| #Shuffles |  |  |  |  |  |
| 2000 |  | 0.84 | 0.86 | 0.84 | 0.84 |
| 1000 |  | 0.82 | 0.83 | 0.84 | 0.87 |
| 500 |  | 0.85 | 0.86 | 0.85 | 0.85 |
| 250 |  | 0.86 | 0.85 | 0.85 | 0.85 |

**Table 5:** STA Significance Test Specificity: Proportion of 100 simulations (per condition) where *none* of the three STA significant tests reported a significant spike-signal relationship, out of all simulations where spike had no effect (Y1, Y4, Y5, Y8).). Here, the only the number of spikes vary in the STA significance calculations (shuffles counts are only used for co-estimated SDO significance). Specificity of the STA significance methods are more variable than sensitivity across simulation. STA significance specificity is lower than for the SDO.

| STA | #Spikes | 250 | 500 | 1000 | 2000 |
| --- | --- | --- | --- | --- | --- |
| #Shuffles |  |  |  |  |  |
| 2000 |  | 0.82 | 0.79 | 0.85 | 0.79 |
| 1000 |  | 0.73 | 0.73 | 0.76 | 0.78 |
| 500 |  | 0.77 | 0.78 | 0.78 | 0.78 |
| 250 |  | 0.78 | 0.79 | 0.77 | 0.78 |

### Identifying when SDOs are needed instead of STAs in Physiological Data

EMG signals (2000Hz) were processed offline using a series of 60 Hz notch (first 8 harmonics), 10 Hz high-pass, and 20-point rectified root-mean-square zero phase filters. Because the distribution of EMG amplitude in our trials was roughly exponential, we log-transformed the processed EMG before assigning states. The resultant assigned and quantized ‘state signal’ varied smoothly and ‘continuously’ (i.e., subsequent discrete states effectively only transitioned to adjacent states when using a 0.5 ms time step), although this was not essential for any of the development and testing here. Under this definition of signal states, we observed signal behavior of the spike-triggered averaged signal waveform (in state space), often varied by signal state at time of spike (**Fig. 6**) (e.g., demonstrated state dependency).

Similarly, and unsurprisingly, the occurrence of spikes was not homogeneous by signal state, as measured by the probability of state at spike, *p*(*x*|*s*). The distribution of state at spike varied with the spiking source tested. Motor activity evoked in our wiping trial paradigm was usually brief and vigorous, with motor unit spikes generally restricted to higher motor states as motoneurons were increasingly recruited during motor behavior. In contrast, recorded interneurons often spontaneously spiked during both motor-quiescent and motor-active epochs of our recording trials. Accordingly, interneuronal spiking events were recorded during both low and high EMG signal states. We thus set about better identifying and predicting these state-dependent effects using SDO approaches.

### Identifying Significant SDO Effects

We used 10 ms intervals (20 points at 2000Hz) before and after (relative to) the spike to generate the *pre-spike p*(*x*_o_) and *post-spike* state distributions *p*(*x*_l_). This interval was selected to examine the immediate post-spike effects on signal behavior (corresponding to 1-3+ chemical synapses). However, arbitrary intervals are possible using the *SDO Analysis Toolkit*.

We proceeded as described in detail in the methods. Stochastic Dynamic Operators were exhaustively generated for every combination of spiking source and signal within the dataset (11 EMG x 283 (likely-redundant) spike trains = 3113 potential SDOs). We defined a ‘significant’ SDO as a spike-triggered SDO matrix which differed from the respective spike-shuffled SDOs for any of the 4 tests of SDO effects as described in the methods, at a P-Value < 0.05. Significant SDOs were used to predict single signal states and post-spike state distributions.

### Example SDO Analyses in spinal data

We compared the STA and SDO methods with the physiological data we collected from spinal frogs. We found that significant spike-triggered SDO matrices generally matched or improved the accuracy of the STA-based method when predicting both single post-spike states and probability distributions. When SDOs were significant, the confidence intervals of the cumulative prediction errors generated using the SDO [H7] vs. STA [H3] matrix hypotheses often had no overlap when using 1000-5000 shuffles (i.e., there were no instances where performance of the STA and SDO were comparable), indicating ‘effect sizes’ in the model lay beyond the observed distribution tails obtained from the bootstrapping (although specific p-Values are ill-defined in these instances). Other hypothesized matrix models had distributions that were more closely positioned and showed distribution overlaps that allowed estimation of probability values. Overall performance of SDOs was thus often much better than other models including classical STA.

Significant spike-triggered SDOs generated from interneurons to EMG signal amplitude often displayed high tuning to signal state and strong state-dependent effects. In the example SDO analysis shown in Figure 9, as indicated by the SDO matrix, a strongly-stereotyped relationship exists between interneuronal spike and signal behavior during high EMG states (**Fig. 9A**). This relationship could also be extracted as the impulse response from the time-domain STA. The ‘arch-like’ rapid rise and fall of EMG signal state, observed in the STIRPD (**Fig. 9B**) is a consequence of using a zero-phase smoothing filter on relatively isolated motor units that were active, and observed to be recruited at high intensities on the aggregate EMG channel. The peak of the filtered signal, and the time of the motor unit occurrence on the channel is observed in the STIRPD at state 20,

**Figure 9:**
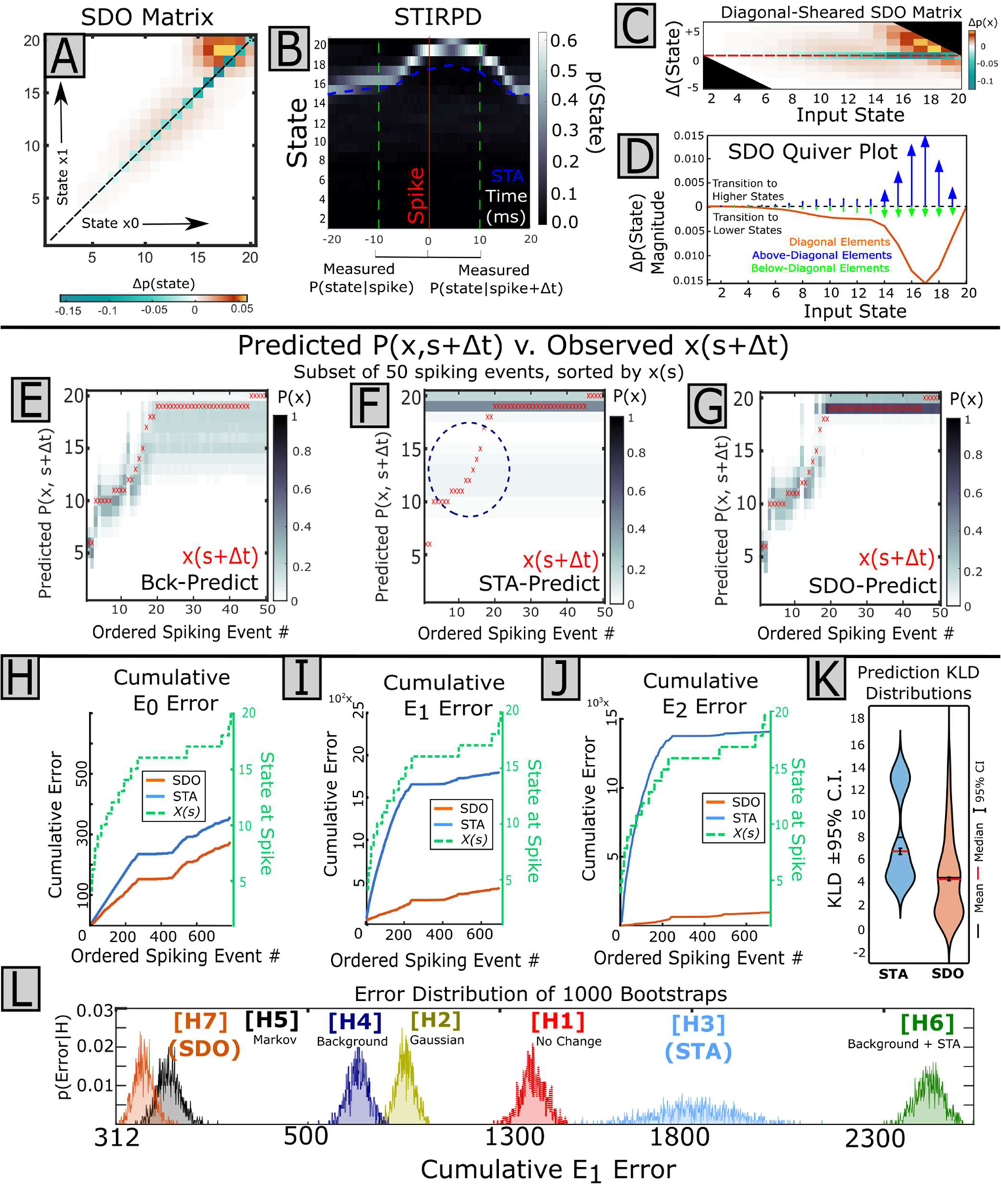
SDO Analysis of a spinal interneuron and EMG amplitude: The spikes from a single spinal interneuron were compared against EMG signal amplitude from the Vastus externus muscle (filtered zero-phase, acausally). **A**) The spike-triggered SDO matrix (here, gaussian smoothed for visualization). Spike-triggered effects are primarily associated with higher states (15-19), with an increased probability of transition towards relatively greater states, as positive elements are above the diagonal. **B)** The extended STIRPD of the EMG signal shows a coarse relationship between spike time and signal state. After spike time, signal state appears to converge on state 18-19 with a probability around 0.6. **C**) The shear SDO shows the positive elements of the matrix are primarily concentrated above the diagonal over states 15-19. This suggests the spike-triggered SDO is consistent with a transition towards higher post-spike states for input states in this region. The slight diagonal orientation of the domains (parallel to the sheared top of the matrix) suggests the post-spike signal state is ‘stepping-up’ to a particular state, rather than broadly increasing state, as first suggested by the STIRPD. **D**) The SDO quiver plot shows the coarse directional effects of the SDO for each input state. Consistent with the shear SDO, the effect of this spike-triggered SDO is to support a transition towards higher states for input states 15-19, indicated by vectors above and below the abscissa pointing upward for these states. **E-G**) For a subset of 50 spiking events, the predicted *post-spike* state distributions were calculated for each spike using the STA or SDO. Each predicted *post-spike* state distribution was represented as a column vector, ordered according to state at spike, and horizontally concatenated into a matrix, displayed here as a grayscale image. The single observed post-spike state for each spike is overlaid as a red x in the respective column. **E)** The background SDO demonstrates state-dependent predictions independent of spike-triggered effects. Here the background SDO is well-suited to predict *post-spike* distributions when in a ‘lower’ initial state, but makes overly-broad predictions at higher states **F)** Here, the STA can predict the post-spike state only over a limited range of experimental data (ordered spiking event 25+). The STA fails to accurately predict *post-spike* state distributions when predicting from a lower pre-spike state (indicated by the blue circle of observed post-spike states not covered by STA-predicted *post-spike* state distributions) but is accurate at higher states. **G)** In contrast, predictions of post-spike state by the SDO are valid over the entirety of the dataset. **H)** When predicting to single states, the SDO reduces both the frequency (E0), **I)** magnitude (E1), and **J**) Sum-squared magnitude (E2) of prediction errors relative to the STA. This predictive accuracy is state dependent: The SDO and STA have equivalent performance at high input states, but the STA significantly underperforms the SDO’s prediction error at lower states, consistent with D. **K)** When predicting *post-spike* distributions, the SDO outperforms the STA (as measured by the Kullback-Leibler Divergence (KLD) between predicted and observed post-spike states). The distribution of KLD, calculated for every observed spiking event, is given as a violin plot. Here, lower values indicate less divergence from the observed distribution, and hence, a better fit. Here the bimodality of the STA violin plot demonstrates the insufficiency of the STA to predict the *post-spike* state distribution for spikes occurring at ‘lower’ states. **L**) Significance of cumulative errors were tested using 1000 bootstraps of E1 errors for all 7 hypotheses. Here, the distributions of SDO-predicted and STA-predicted errors do not overlap; P-values are arbitrarily small.

∼6 ms *after* the spike for the recorded interneuron. That is, the interneuron in Figure 9 consistently fires before spiking in the vastus externus (VE) EMG channel. However, this relationship between interneuron and motor unit in the aggregate EMG is *state-dependent*: While the VE is in a signal state >14, a spike from this interneuron reliably predicts an increase in VE signal state (consistent with recruiting additional individual motor units). However, when the VE is in states 1-14, the spiking of the recorded interneuron no longer predicts a change of signal state. The SDO is tuned to this relationship, and only demonstrates an effect at these higher states (**Fig 9C,D**). In contrast, the simple classical time-domain STA averages over both conditions. When predicting post-spike states and distributions, pre-spike state-dependency is important: The H3 STA matrix is accurate in predicting when signal state at spike is high (e.g., states 18-20), but is inaccurate when signal state at spike is low (**Fig. 9F**). In contrast, the SDO-based prediction captures this state-dependency, matching the H3 STA prediction accuracy for high states, also greatly improving predictions for the lower states (**Fig. 9G**). The state-dependency background SDO generates reasonable predictions, independent of spike-triggered effects (**Fig. 9E**). If cumulative prediction errors are quantified over all occurrences of spike, the SDO significantly, and greatly, reduces both the frequency (e0; STA = 356.95^k^, SDO = 287.8^l^, Cohen’s D =-10.4) and the magnitude (e1; STA = 1816.8^m^, SDO=487.5^n^, Cohen’s D=-15.5) of the associated cumulative prediction errors relative to the STA (**Fig. 9H-J**). The SDO and STIRPD thus capture a richer description of the spike-triggered signal behavior compared to the classical STA alone (**Fig. 3**). Predicted post-spike *distributions* deviate less from observed distributions using the SDO rather than the H3 STA, measured using a smaller Kullback-Leibler Divergence (KLD) (**Fig. 9K**). (Note that the empirical distributions, **Fig. 9L**, lower, of these different bootstrapped cumulative errors do not overlap, and hence p-Values cannot be directly estimated, (i.e., p-values would lie in long tails of the distributions that were never visited or captured in the bootstrapping of distributions, would be significant, but the specific p-Values are ill-defined in these instances). If use of the classical STA is desired, the SDO and STIRPD can here inform about which spiking events should be used to estimate the STA impulse response, and in what conditions it applies. Taken together, the combination of interneuron state tuning and spike-triggered effects can form the basis for the analysis of potential dynamical neural controls of motor behavior.

In contrast to interneurons, vertebrate single motor units should be causally linked only to a single muscle’s aggregate EMG and force. Nonetheless, intermuscular coordination may theoretically permit significant *correlations* between single motor units in one muscle relative to muscle activity in another. SMUs collected in our dataset demonstrated somewhat stereotyped motor tuning to signal behavior over the experimental interval. Likely unsurprisingly, tuning of aggregate EMG and motor unit was strongest for the parent muscle, although more complicated relationships could also be observed for synergist muscles (**Fig. 10**). In the example SDO analysis, the SDO matrix shows two regions of spike-triggered effects (**Fig. 10A,C,D)**. Here, aggregate EMG signal behavior collected in a synergist muscle around SMU spike times visibly separates based on signal state at SMU spike within the STIRPD (**Fig. 10B**), and *p(state|spike)* and thus resembles a bimodal distribution. Just as the mean of a bimodal distribution may be a suboptimal description of the distribution, the STA calculation averages over this observed separation in signal behavior when predicting post-spike state in the synergist (**Fig. 10F**). Consequently, the H3 STA matrix predicts overly broad *post-spike* distributions, while the SDO predicted *post-spike* distributions that better resembled the observed distributions (**Fig. 10G**). This improvement of the SDO over the H3 STA is similarly observed for predictions to most likely (singular) post-spike state (**Fig. 10I-K**). In this case, SDO methods resulted in significantly, and greatly, lower cumulative e1 (magnitude) error rate compared to the STA (e1; STA = 534.8°, SDO =167.9^p^, Cohen’s D= −20.2). As diagrammed in **Fig. 10L**, empirical distributions of these bootstrapped cumulative errors do not overlap, (i.e., p-values would lie in long tails of the distributions that were never visited or captured in the bootstrapping process, would be significant, but specific p-Values are ill-defined in these instances, being beyond the observed distribution tails’ ranges). Similarly, predicted post-spike *distributions* deviate less from observed distributions using the SDO rather than the H3 STA, measured using a smaller KLD (**Fig. 10K**).

**Figure 10:**
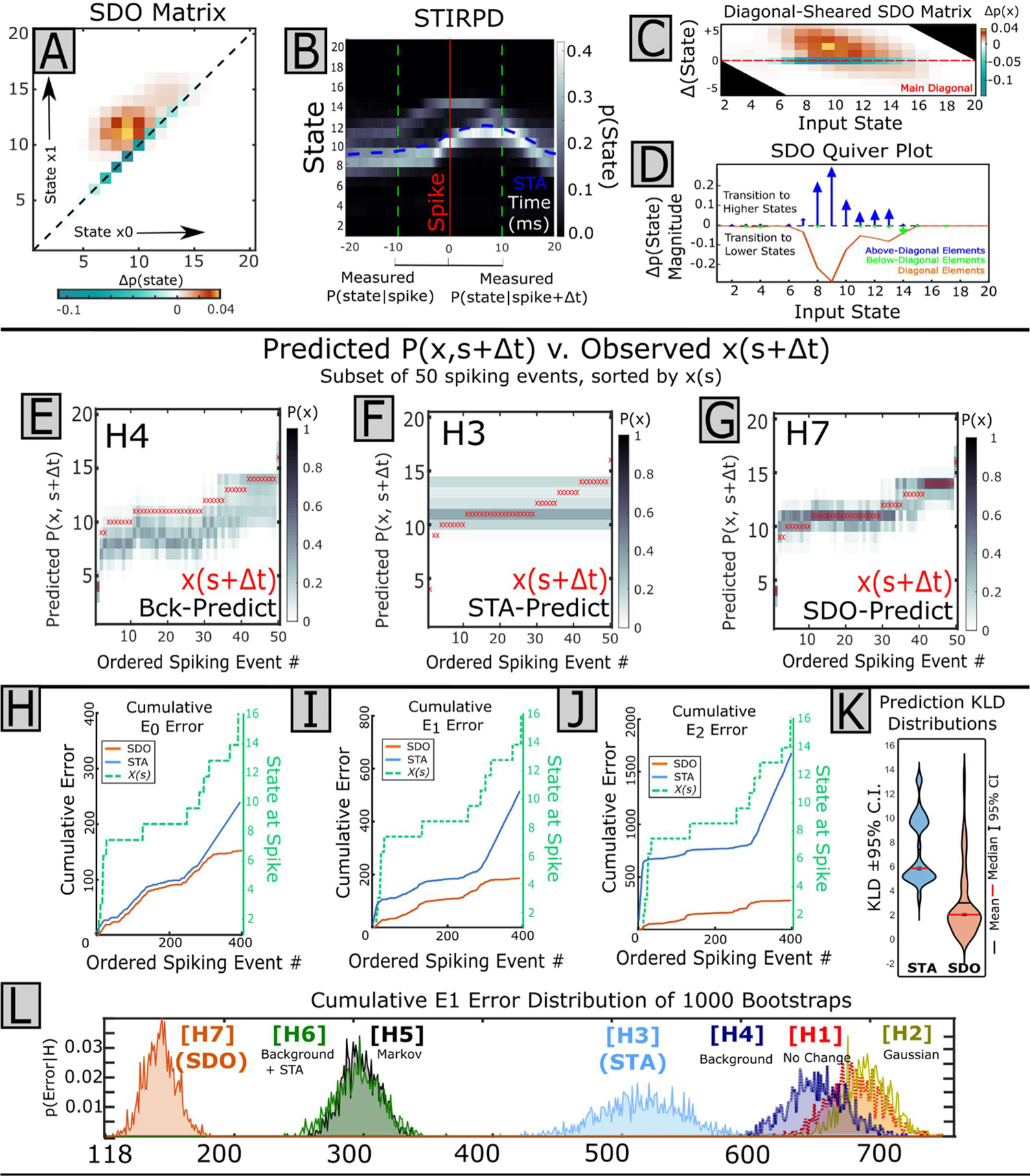
SDO Analysis of a Single Motor Unit and Synergist Muscle EMG: The spike train of a single motor unit (SMU) in the vastus externus muscle was compared against EMG signal amplitude of the biceps femoris (as in **Figure 3**). **A**) The SDO matrix. Effects are localized in two regions, about State 8-10 and 12-14. **B)** The extended STIRPD of the EMG shows this SMU is tuned to two different states of EMG signal amplitude, as indicated by the bimodal behavior of p(state|spike). In the upper ‘arc’, at state 14, the *post-spike* state distribution appears mostly symmetrical to the pre-spike state. However, in the lower arc (state at spike = 10), the post-spike states are increased relative to spike. **C**) The shear SDO shows the positive elements of the matrix are primarily concentrated above the diagonal over states 8-12, corresponding to the lower ‘arc’, but minimal effects outside this region. **D**) Similarly, the quiver SDO demonstrates coarse directional bias towards higher post-spike states for input states 8-12, corresponding to the ‘lower arc’ on the STIRPD, but minimal effects for input states 13-16, consistent with minimal change to the ‘upper arc’ of the STIRPD. **E-G**) For a subset of 50 spiking events, the predicted post-spike state distribution was calculated for each spike using the STA or SDO. Each predicted *post-spike* state distribution was represented as a column vector, ordered according to state at spike, and horizontally concatenated into a matrix, displayed as a grayscale image. The single observed post-spike state for each spike is overlaid as a red x in the respective column. **E**) Predictions from the background SDO are state-dependent although inadequately capture spike effects. **F)** The bimodal *post-spike* distribution predicted from the STA suboptimally predicts the observed post-spike state, while **G)** the SDO predicted distribution of post-spike state more tightly fits the observed post-spike state, across all signal states. Thus, the SDO provides a more reliable method of predicting signal behavior. **H-J**) When predicting single post-spike states, the rate of error accumulation for different hypotheses depends on state. Post-spike signal state was predicted for every spiking event, for all hypotheses, and the error between each event-wise prediction was accumulated for all spikes. Spiking events were sorted by state at time of spike to uncover state-dependent error rates. Here the rate of accumulation for the **H)** frequency (E0), **I)** magnitude (E1), and **J)** Sum-squared magnitude (E2) error are comparable for states 7-10 for the STA and SDO as indicated by the parallel traces of the cumulative error over this region), but the STA performs poorly at lower and higher states, whereas the SDO maintains prediction accuracy over all states. **K**) The similarity between each predicted and observed post-spike distribution was assessed as the Kullback-Leibler Divergence (KLD). The distribution of the KLD over all spiking events is displayed as a violin plot. Predicted distributions using the SDO resulted in a better fit than the STA. **L**) Significance of cumulative errors were tested using 1000 bootstraps of E1 errors (For all 7 hypotheses, below). As suggested by **E** and **F**, the STA results in a better prediction than the background SDO but worse than the spike-triggered SDO. Here, the distributions of SDO-predicted and STA-predicted errors do not overlap; P-values are arbitrarily small.

This improvement in prediction is consistent with SDO methods capturing a richer description of the spike-related state-dependent effects relative to the classical STA. While increasing the number of parameters within a model tends to improve the fit to observed data, the SDO and probability distributions (and H3 STA), as utilized here, are treated as discrete representations of fundamentally smooth and continuous objects. Hence, from an Akaike or Bayes Information criterion on model complexity the exact number of parameters, and degrees of freedom (DOF), vary with chosen quantization of state (e.g., a gaussian distribution measured within 100 binned intervals may possess only 2 fundamental degrees of freedom.) However, in predicting *p*(*x*_l_|*x*_o_) instead of *p*(*x*_l_), the SDO likely always represents more DOF than the classical STA. Nonetheless, the information gain in using the SDO-predicted post-spike distribution to represent the observed distribution (as indicated by the KLD, Fig. 9K,10K) is greater than for the STA-predictions, the efficiency of this gain depends on SDO matrix dimension choices and data available.

### Classification of SDO types and Motifs

When the state-quantized signal is smoothly varying and state distributions are drawn using a short interval, non-zero elements of the SDO are usually concentrated near the matrix diagonal. Accordingly, the SDO is a sparsely sampled matrix, with zero-value elements where state transitions are not observed and non-zero elements reflecting the extent of *P(state|spike)* over all spiking events. Positive elements of the SDO indicate increased probability of the state transition (*j→ i*), while negative elements indicate decreased probability of transition (j→ i). This permits the localization of positive and negative domains in the SDO to be descriptors of the correlated ‘actions’ of a spike on signal behavior (**Fig. 11**).

**Figure 11:**
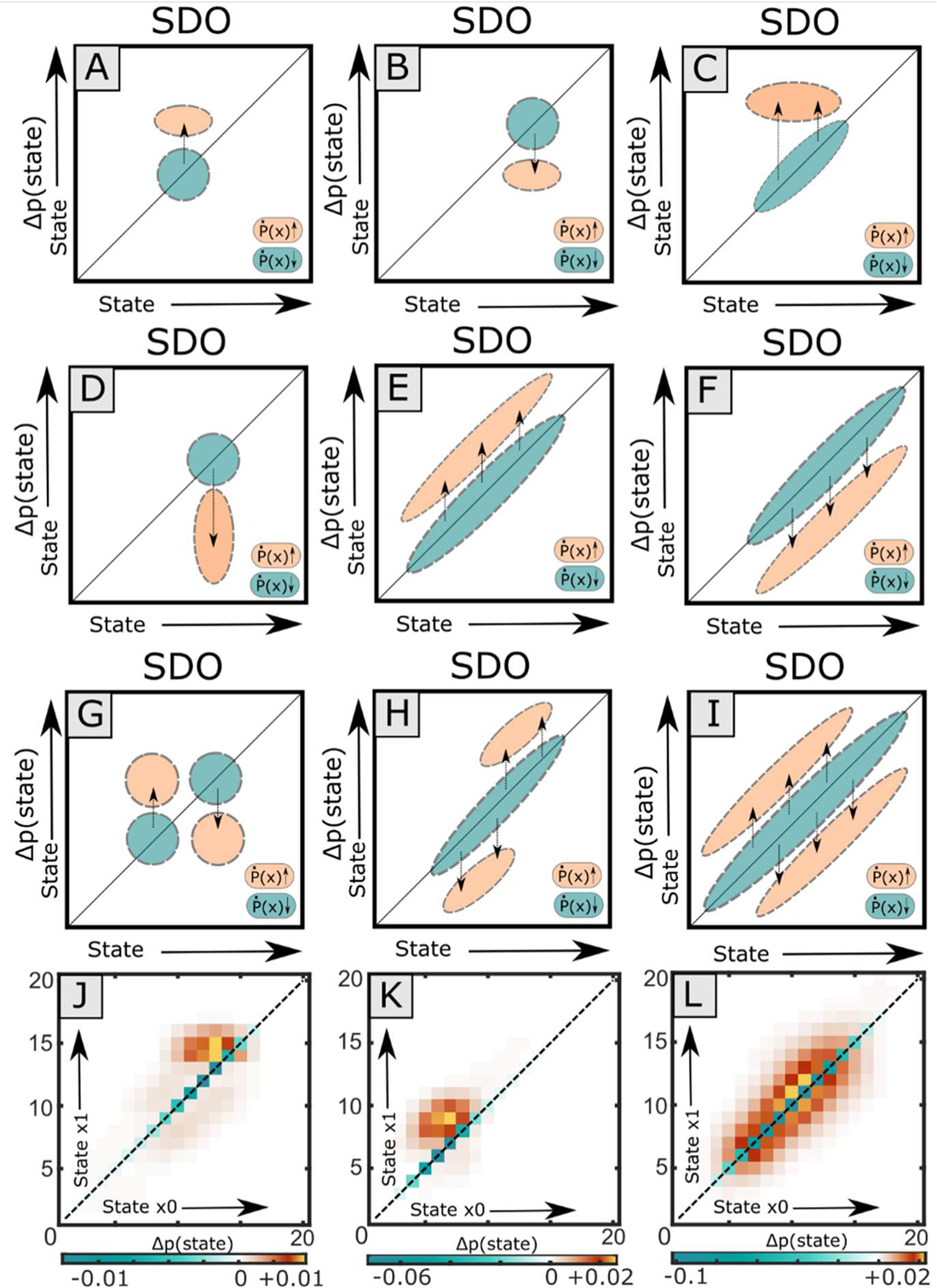
The effect of the SDO in transforming the pre-spike distribution into the post-spike distributions can be characterized and crudely classified from the localization of Δp(x) in the SDO array. SDO features may have effects arranged along the matrix diagonal or display horizontal or vertical banding. Topologies may be combined to accumulate effects in a state-dependent fashion. The direction of the dashed arrows indicates the general direction that pre-spike distribution of state is modified by the SDO. **A)** *Step Up Operator*. **B**) *Step Down Operator*. **C**) *Convergence Operator* (up). **D**) *Divergence Operator* (down). **E**) *Increase Operator*. **F**) *Decrease Operator*. **G**) *Stabilize State Operator*. **H**) *Destabilize Operator.* **I**) *Diffusion Operator*. **J**) An example ‘Convergence’ SDO. **K**) An example ‘Increase’ SDO. **L**) An example ‘Diffusion’ SDO.

The state-dependent changes in *post-spike* distribution enacted by an SDO could be inferred by viewing the matrix directly. We identified 9 classes of simple features or ‘motifs’ which represented specific types of changes observed in the data. These motifs are summarized in Figure 11 for sparse matrices. Figure 11 shows important examples of simple sparse SDOs forming elementary ‘motifs’. When evaluating motifs, both the normalized and parameterized SDOs should be considered: The normalized SDO displays spike-triggered ‘effects’ while the parameterized SDO weights these effects by the frequency of occurrence. This summary helps illustrate how SDOs can be intuited and interpreted when the SDO matrix is sparse. These SDO motifs can be classified as below.

#### ‘Step-Up/Down’ Operator: (**Fig. 11A, B**)

These elementary SDOs are characterized by a restricted negative domain on the matrix diagonal, with a similarly-restricted positive domain above (**Fig. 11A**) or below (**Fig. 11B**) the diagonal. This restriction limits the number of states modified by the SDO, and generally reflects motor tuning for the spike-signal relationship. The *Step-Up* and *Step-Down* operators decrease the probability of maintaining state and increase the probability of transitioning to a narrow band of post-spike states, from a limited band of input pre-spike states. This elementary operator was the most common from single motor units to EMG which demonstrated significant motor unit tuning.

#### ‘Convergence’ Operator (**Fig. 11C**, and 11J from spinal data)

This elementary SDO is characterized by a *horizontal* band. The *Convergence* SDO increases the probability of transitioning *to* a restricted band of states in the post-spike distribution from a larger band of input (*pre-spike*) states. (Note that this domain may occur above or below the matrix diagonal.) This SDO usually corresponds with a decreased variance of the post-spike distribution. This operator was commonly extracted in the normalized SDO matrix when there was a strong reversion to a common post-spike state distribution (i.e., converging on a common state distribution). **‘Divergence’ Operator** (**Fig. 11D**):

This elementary SDO is characterized by a *vertical* band. The function of *Divergence* SDO is opposite of the *Convergence* SDO: pre-spike input states within a limited band (defined by the horizontal breadth of this motif) are distributed to a broader output band of states (defined by the vertical breadth of the positive domain of this motif), which may be above, below, or on either side of the matrix diagonal. This motif may emerge if the range of states in the pre-spike distribution is limited (e.g., spike state tuning), but the range of states in the *post-spike* distribution of states is not as tightly controlled. This bidirectional increase in signal dynamic range can also occur when the spike impulse response is multimodal (such as a single motor unit action potential,), and the signal fluctuates stochastically about a mean value (such as unrectified EMG or the simulated white noise process).

#### ‘Increase/Decrease’ Operator (**Fig. 11E, F, and** 11K from spinal data)

These elementary operators are characterized by a positive *diagonal* band either above (**Fig. 11E**) or below (**Fig. 11F**) the main diagonal. Because positive domains run parallel to the matrix diagonal, the effect of the *Increase* or *Decrease* operator is broad, describing a shift of the post-spike distribution towards increased or decreased signal state. If these bands are adjacent to the main diagonal, these operators permit a smooth drift of the *post-spike* distribution. Crudely, these would correspond to excitatory or inhibitory synaptic effects, respectively, if observed neuron-to-neuron or neuron-to-EMG. This SDO motif maps reasonably well onto the classical STA picture for the states of origin. Indeed, in simulated data, the H3 matrix (STA) produced this SDO when there was a clear and consistent post-spike facilitation or inhibition of signal amplitude.

#### ‘Stabilize’ Operator (**Fig. 11G**)

This SDO resembles the co-occurrence of the *Step-Up* and *Step-Down* operator acting over a clustered band of states. When the input *pre-spike* distribution trends towards lower states, the *Stabilize* operator increases probability of transition towards *higher* states (‘step up’). Simultaneously, when the input *pre-spike* distribution trends towards higher states, the *Stabilize* operator increases the probability of transition towards *lower* states (‘step down’), the net result stabilizes the probability of state in the center of the motif. This is equivalent to a negative feedback controller around the setpoint state at the center of the motif and was indeed recovered from both the background and spike-triggered SDOs from stabilizing dynamical systems (e.g., Y4, Y7).

#### ‘Destabilize’ Operator (**Fig. 11H**)

This SDO resembles the co-occurrence of an *Increase* and *Decrease* Operator concatenated along the diagonal. The function of the *Destabilize* operator is inverse of the *Stabilize* operator: Lower input states are pushed towards lower output states, and higher input states towards higher output states. Input states near the middle of the *Destabilize* operator will be ‘destabilized’ and may be pushed towards a bimodal post-spike distribution, away from the initial state.

#### ‘Diffusion’ Operator (**Fig. 11I**, and 11L from spinal data)

This SDO is characterized by a negative region along the diagonal, with positive diagonal bands both above and below the diagonal. The function of the *Diffusion* SDO is to broaden the distribution of state (i.e., increase the variance of the *post-spike* probability distribution), similar to diffusion in a fluid. If the magnitude of *Δp(x)* above and below the matrix diagonal is comparable within a column of the SDO matrix, the increase in variance due to this operator does not necessarily change the mean of the *post-spike* distribution. The *Diffusion* Operator was the most common SDO in our physiological datasets when describing background signal dynamics with shuffled timestamps. Examples of 3 such motifs from our physiological data are shown in **Fig. 11J-L**.

## DISCUSSION

We have demonstrated that SDO methods capture dynamics and spike-correlated information better than the classical STA in both real and simulated data sets. While we have demonstrated that the SDO is a useful method, the extra considerations it offers may not be necessary for all conditions. In simulation, the tested STA methods of significance were sensitive to both consistent (Y3) and random (Y2) spike-triggered impulse responses within white noise, and in signals on a dynamic background (Y6). Random effects may be important, but are not part of classical circuit-breaking STA identification. The SDO description makes such stochastic effects explicit. However, classical STA is robust and computationally simple: Signal is collected around spiking events and averaged together. The extracted waveform can serve as a template for the correlated ‘effect’ of the STA which can be measured as the change in this average signal. This is particularly useful for reverse correlation. Similarly, because the average signal envelope is calculated independently for each aligned time bin in the STA, the duration of the pre-spike and post-spike windows generally do not affect significance of the STA. However, the STA was incapable of consistently assigning significance when spike-triggered effects were dynamic (Y7). Further, SDO-based tests of significance were both more sensitive and specific than those of the classical STA for identifying significant state-dependent relationships between spike and signal across all simulated data. This is likely a function of the state-dependent operations and bootstrapping procedures standard to the SDO method. When applied to physiological data, our results show that SDO analysis provided superior, more nuanced descriptions of spike-correlated signal behavior than the classic STA impulse response when complex interactions were present. This held true for both spinal interneurons and single motor units’ correlations to homonymous or synergist EMGs. The STA estimates the spike-triggered response amplitude using the means of the distribution of observed amplitudes across all spikes. Thus, the STA depends on assumptions of mean and normality as optimal descriptors of the response distribution. The example data here show these assumptions are violated by multimodal distributions of output state (e.g., STIRPD **Fig 10B**), and state-dependent output behavior (as in **Fig 9B**). While these factors may be handled with additional explicit consideration with the STA, because SDOs are probabilistic descriptors, SDO methods provide tools that straightforwardly capture these features of neural data automatically.

SDO methods measure changes in *distributions* of signal state rather than the average amplitude of the signal at every time index. This method trades sensitivity in the time domain for sensitivity to signal variance. When there are strong relationships between signal and time, such as motor units to the homonymous muscle in the data analyzed here, the accumulation of state into *pre-spike* and *post-spike* distributions for estimating the SDO may compress time-domain information depending on binning choices, that can be observed in an STA signal envelope. In these cases, the STIRPD with appropriate binning provides a reasonable compromise, behaving as a quantized STA.

### Caveats of State for Stochastic Dynamic Operators

Because the SDO describes changes in state, the definition of state from original signal amplitude is important. The number of bins, selection of signal levels, filtering, and transformation of the signals (e.g., log-transform) will all change the representation of the state space time series. The definition of state (i.e., the quantization scheme) must be sufficiently sensitive to capture changes in signal state distribution needed to detect significant correlations with the SDO: If signal amplitude before and after spike is assigned to the same state, there will be no measured change in *pre-spike* and *post-spike* distributions (i.e., the SDO matrix will resemble our null hypothesis H1). This sensitivity can be established *a priori* because state is often derived from scaled amplitude ranges of signals with real units (e.g., for a signal with a dynamic range of 10 mv, 20 equal-width binned states are sensitive to effects >|0.5 mv| of original signal). State bin width determines the minimum detectable effect in state space. Explicit, intentional, determination of dynamic range will likely be necessary for detecting significant effects with the SDO. As the number of states increases, sampling of state-to-state transitions observed within a finite dataset necessarily decreases. Hence, state numbers used in SDO analyses represents a tradeoff between resolution, information, sampling theory, experimental constraints, and resulting statistical power. In our results, 20 states were sufficient to detect significant correlations between spikes and signal behavior, without requiring an inordinate number of spikes.

Similarly, temporal relationships between spikes and distributions constrain spike-triggered SDOs. Shorter time windows are clearly more sensitive to high-frequency (quantized) signal content than longer time windows which effectively filter the high frequency, as with any choice of binning intervals. Latency in determining *when* to define ‘pre-spike’ and ‘post-spike’ intervals, is similarly important (*i.e.*, is effect of spike near-immediate, or delayed and by how much?). SDO predictions thus capture spike effects strictly in context of experimenter signal processing and analysis choices. However, analogous decisions also affect classical STAs (although these are not always explicit or considered). Spike-triggered SDO effects are best described using brief time windows when drawing probability distributions near spike, and when background stochastic signal is stationary over the time course of the windowed signal. In the examples here, we presented one latency and binning strategy for purposes of demonstration. When tested in simulation, this combination permitted the SDO to be both more sensitive and specific than the STA using only 250 spikes and 1000 shuffles. To increase utility, we permit these parameter choices to be user-defined within the *SDO Analysis Toolkit*.

Predictions of both *post-spike* state distributions and singular (‘best’) states were compared between STA (H3) and SDO approaches. Experimentally, most arbitrary combinations of spiking source and signal did not produce significant SDOs (or STAs) in physiological data. Non-significant SDOs (using our 5 measures of matrix significance), usually resembled *Diffusion* Operators (hypothesis H2, and Figure 11 panel I, and L), enacting an unbiased passive increase in signal variance over time. Diffusion operator SDOs could, in principle, be statistically significant, but were not in our data. Nonetheless, spike-triggered SDOs which were significant generated superior predictions relative to H3 STA. However, all 7 model hypotheses were implemented as matrices which satisfy the definition of an SDO, albeit generated by different strategies. Hence, even if the spike-triggered SDO matrix (H7) estimation is suboptimal compared to the other prediction model hypotheses, the model of best fit may still be interpreted using SDO methods.

### Interpretations of SDOs and State Tuning in Examined Spinal Data

In vertebrates, single neurons usually act as part of larger organizational units. It is thus not unreasonable to assume the effects of a neuron will depend on the activity of covarying units and system state. Spinal interneurons must, to some degree, be both sensitive to ongoing motor state and capable of influencing motor state (e.g., AuYong, et al., 2011A). Classical STA can show clear interneuron effects (e.g., Hart and Giszter, 2010, Takei et al. 2017). Similarly, significant SDOs in our data were also extracted when signals showed such statistically consistent behavior in the post-spike interval, as visualized on the STIRPD (e.g., **Fig. 9,10**). However, the correlation between interneurons and motoneurons to EMG were also sensitive to pre-spike signal state in our data. SDO methods could detect and capture these pre-spike dynamics.

In our data, unsurprisingly, significant SDOs were often observed between single motor units (SMU) and homonymous muscle EMG. This is consistent with the motor unit summation constructing the recorded EMG signal following the size principle (Henneman, et al, 1965). However, significant SDOs also existed in our data for the same spiking SMU against EMG signals across multiple muscles, with similar motor tuning. For example, the stereotyped ‘arcing’ observed in distributions for the biceps femoris muscle (STIRPD **Fig. 10B**) corresponded to the likely spiking of correlated motor units, recorded in the aggregate EMG implanted into the biceps muscle. The reliable co-recruitment of motor activity in biceps’ aggregate EMG accompanied the single motor unit in the vastus externus, recorded with fine-wire braided probes, is consistent with synergy-based control of motor pools, and also some features of human data reports (Laine, et al, 2015) (Del Vecchio, et al, 2022) (Hug, et al, 2023) (Negro et al.,2016).

Vertebrate SMUs are confined to a single muscle: SMUs thus cannot be directly causal to heteronymous muscle EMG. However, in synergy premotor drives and in the spinal reflexes, activity of single motor units may be tuned to the state not only of the homonymous muscle, but also synergists and antagonists. Insofar as muscles have correlated coactivation at (fast) time scales examined here, synergist motor unit spikes and EMG will correlate during observed co-contractions. Common synergist premotor drives could directly induce such correlations. Such correlations may be state-dependent: Synchronization between motor units is more frequently observed at low-force levels than during high-forces (e.g., Basmajian and De Luca, 1985, Semmler, et al, 2000). SDO-based methods can identify and estimate stochastic and state-dependent effects, which may provide additional predictive power in analysis of these correlations.

### SDO Sensitivity and Caveats

For use of the STA and SDO as explored here, we treated spiking events and spike-signal correlates as independent, with the spike impulse response as linearly estimated across samples (although the SDO permits for state-dependency in response). In circumstances where signal likely facilitates spike (such as stimulus-triggered averaging in the visual system), STA of the stimulus in the pre-spike interval can identify the average filter (or static combination of filters) which are the maximum likelihood estimation of the signal facilitating a response. In these cases, deviations of the neuron from the Linear-Nonlinear-Poisson (LNP) model implicit to the STA become particularly important: Estimating the average post-spike facilitation of a bursting neuron may be straightforward but estimating the stimulus triggering the burst (if indeed it is a burst) may not. In these instances, both the classical STA and SDO might sometimes produce spurious correlates. Alternative estimations of the reverse-correlated STA have been proposed to manage deviations of the neuron from the LNP model (e.g., Paninski et al., 2004), but these have not been implemented for the use of the SDO for forward-correlation between spike and signal as presented here.

Neither SDOs nor classical STAs can directly infer causality between a spiking process and a signal. Determining causality between observed processes requires explicit experimental manipulations (e.g., driving the neuron). Nonetheless, correlative models are useful predictive tools, and can indicate the need for further causal tests. (Indeed, in neural encoder/decoder frameworks, the ‘function’ of a neuron may be divorced from its capacity to reflect or predict network state.) Here, SDO analysis can augment other correlative tools such as blind source separation methods (e.g., independent components analysis and non-negative matrix factorization). SDO frameworks can integrate correlation patterns and recordings of interneurons, as observed in our data with interneurons predicting signal states across multiple muscles. When neurons are known, or modelled, as differentially contributing to motor behavior by neural configuration (e.g., segregation of rhythm and pattern in locomotion (McCrea and Rybak, 2008; Danner, et al, 2017), and spinal ‘state’ (Ausborn, et al, 2018)), SDOs methods may provide exploratory tools. As far as different SDO motifs capture significant state-dependent spiking behaviors within the data, different classes of neurons should generate different SDO motifs: SDO matrices are thus likely useful descriptive classifiers.

The SDO framework can be extended. We demonstrated SDO analyses using single-channel EMG time series data and stochastic simulated data, although more complex applications are likely to be straightforward. In the depiction of the SDO presented here, ‘state’ is defined using amplitude of the same signal correlated with spike. The SDO may be extended to incorporate additional orthogonal dimensions of state by evaluating changes to joint probability of the form Δ*p*(*x*_l_|*x*_o_, … θ), where θ can represent any number of independent quantizable parameters. (Although in this instance, the dimensionality of the SDO matrix will grow by the number of discrete variables and may no longer be interpretable as a 2D matrix). SDOs may thus be scaled as necessary to address the experimental question.

SDO analysis may thus be applied to combined or derived signals (*e.g.,* activations of independent or principal components, rate processes, neural ensemble-derived dimensionality reductions). For example, spiking events may be correlated with projection component features obtained from higher-dimensional datasets (see Smith and Brown, 2003; Churchland, et al., 2008; Auyong, et al., 2011B; Raposo, et al., 2014; Lindén, et al, 2022; McMahon, et al., 2022; Thura, et al., 2022). The extension of SDOs to oscillatory processes via the definition of a stochastic phase variable may be of particular use for linking single-unit effects with population dynamics (e.g., neurons may be sensitive to network phase, or evoke phase-dependent effects), but will require additional validation than has been presented here. On these frontiers, SDOs provide a new means to visualize and explore predictive influences of specific spike trains in these dynamics and based on state variables.

Our SDO methods described here provide a framework for experimentalists to explore state-dependent and probabilistic relationships within data, and to test generative parameters within the *SDO Analysis Toolkit*. A repertoire of statistical tests is used to identify significant SDO matrices, and to test SDO-based predictions. SDO matrices are tested for significance using Monte-Carlo bootstrapping using our five tests of matrix composition. Predictions of signal behavior are tested using our seven matrix hypotheses, including the SDO and the STA, to determine the model of best-fit in observed data to both single state and distributions. While we cannot provide an exhaustive survey of potential SDO topologies, Figure 11 provided a rudimentary classification to interpret SDO ‘function’ within recovered SDOs. We believe the combination of visual intuition with mathematical foundations that provided by SDO analysis support more precise and nuanced analyses of spike effect relative to the classical STA alone.

## Conclusions

Here we introduced and demonstrated Stochastic Dynamic Operators as useful analysis tools in both physiological and simulated data. SDOs extend classical STA into probabilistic and state-dependent domains, behaving as a state-dependent spike-triggered average which may better capture spike correlations and potentially causal effects. The SDO offers improved sensitivity and specificity for identifying significant relationships between the spike and signal relative to the classical STA. The utility and function of SDO methods can be intuited through motifs on the SDO matrix, the shear and quiver SDO visualizations, and the spike-triggered impulse response probability distributions (STIRPDs). Together, these methods capture classic STAs and graphically reveal significant and state-dependent stochastic behaviors. SDO analysis is thus a novel method for both qualitative and quantitative data interpretation. We demonstrated the utility of SDOs as a flexible tool for describing relationships between spinal interneurons, single motor units, and aggregate muscle activations. We showed that using SDOs and STIRPDS in place of the classical STA can better characterize post-spike effects, account for non-correlated background dynamics, improve sensitivity and specificity of detecting significant spike-signal correlations, and significantly increase prediction accuracy. SDOs provide a statistical model of neural actions, are easily obtained from existing data, and can be empirically tested for statistical significance. The SDO may be readily built with data collected on multiple time scales and experiment types, including stimulation and interventions, across varying contexts, which far exceed applications here in simulated and spinal motor control data. In summary, SDO analysis is a powerful but accessible method, and we anticipate both the framework and *SDO Analysis Toolkit* here to augment the general Neuroscience toolbox.

## Code and Accessibility

The SDO analysis and data figures were produced using the *SDO Analysis Toolkit*. The code/software described in the paper is freely available online at https://github.com/GiszterLab/SdoAnalysisToolkit

---

**Significance Tables.** Types of tests and measures of confidence intervals used. The row label corresponds to the superscript of the matching test.

| Data Structure |  | Type of Test | Confidence Intervals |
| --- | --- | --- | --- |
| a | Normal | T-Test (1-tailed) | 39.31 ± 2.425 |
| b | Normal | T-Test (1-tailed) | 99.81 ± 0.198 |
| c | Normal | T-Test (1-tailed) | 27.69 ± 1.517 |
| d | Normal | T-Test (1-tailed) | 15 ± 3.307 |
| e | Normal | T-Test (1-tailed) | 22.31 ± 1.637 |
| f | Normal | T-Test (1-tailed) | 17.88 ± 4.111 |
| g | Normal | T-Test (1-tailed) | 17.06 ± 2.722 |
| h | Normal | T-Test (1-tailed) | 2.688 ± 1.635 |
| i | Normal | T-Test (1-tailed) | 19.09 ± 2.622 |
| j | Normal | T-Test (1-tailed) | 15.36 ± 5.780 |
| k | Bootstrapped Normal | T-Test (1-tailed) | 357 ± 0.810 |
| l | Bootstrapped Normal | T-Test (1-tailed) | 287.9 ± 0.775 |
| m | Bootstrapped Normal | T-Test (1-tailed) | 1816.8 ± 5.863 |
| n | Bootstrapped Normal | T-Test (1-tailed) | 487.5 ± 1.964 |
| o | Bootstrapped Normal | T-Test (1-tailed) | 534.8 ± 2.003 |
| P | Bootstrapped Normal | T-Test (1-tailed) | 167.9 ± 0.781 |

## Notes

https://github.com/GiszterLab

### Competing Interest Statement

The authors have declared no competing interest.

## References

Ausborn J, Snyder AC, Shevtsova NA, Rybak IA, Rubin JE (2018) State-dependent rhythmogenesis and frequency control in a half-center locomotor CPG. J Neurophysiol 119(1):96–117

AuYong N, Ollivier-Lanvin K, Lemay MA (2011) Preferred locomotor phase of activity of lumbar interneurons during air-stepping in subchronic spinal cats. J Neurophysiol 105(3):1011–22.

AuYong N, Ollivier-Lanvin K, Lemay MA (2011) Population spatiotemporal dynamics of spinal intermediate zone interneurons during air-stepping in adult spinal cats. J Neurophysiol 106(4):1943–53

Basmajian JV, De Luca CJ (1985) Muscles Alive: Their functions revealed by Electromyography. Baltimore, MD: Williams and Wilkins.

Bizzi E, Mussa-Ivaldi FA, Giszter S (1991) Computations underlying the execution of movement: a biological perspective. Science 253(5017):287–291.

Bokil H, Andrews P, Kulkarni JE, Mehta S, Mitra PP (2010). Chronux: a platform for analyzing neural signals. J Neurosci Methods. 2010 Sep 30;192(1):146–51.

Buys EJ, Lemon RN, Mantel GW, Muir RB (1986) Selective facilitation of different hand muscles by single corticospinal neurons in the conscious monkey. J Physiol 381:529–49.

Cheney PD, (1980) Response of rubromotoneuronal cells identified by spike-triggered averaging of EMG activity in awake monkeys. Neurosci Lett 17(1-2):137–42.

Cheney PD, Fetz EE (1985) Comparable patterns of muscle facilitation evoked by individual corticomotoneuronal (CM) cells and by single intracortical microstimuli in primates: evidence for functional groups of CM cells. J Neurophysiol 53(3):786–804.

Churchland M, Yu BM, Sahani M, Shenoy KV (2008) Techniques for extracting single-trial activity patterns from large-scale neural recordings. Curr Opin Neurbiol 17(5):609–618.

Cohen J (1988) Statistical Power Analysis for the Behavioral Sciences. Mahwah NJ: Lawrence Erlbaum Associates.

Danner SM, Shevtsova NA, Frigon A, Rybak IA (2017) Computational modeling of spinal circuits controlling limb coordination and gaits in quadrupeds. Elife 6:e31050.

Davidson AG, O’Dell R, Chan, V, Schieber MH (2007). Comparing effects in spike-triggered averages of rectified EMG across different behaviors. J Neurosci Methods, 163(2), 283–294.

Del Vecchio A, Germer C, Kinfe T, Nuccio S, Hug F, Eskofier B, Farina D, Enoka R, (2022) Common synaptic inputs are not distributed homogeneously among the motor neurons that innervate synergistic muscles. *bioRxiv* 477379.

Devanandan MS, Eccles RM, Yokota T (1964) Presynaptic Inhibition induced by muscle stretch. Nature 204:996–8.

Figueiredo MAT, Jain, AK (2002) Unsupervised Learning of Finite Mixture Models, IEEE Trans Pattern Anal Mach Intell 24:381–396.

Fetz EE, Cheney PD (1980) Post-spike facilitation of forelimb muscle activity by primate corticomotoneuronal cells. J Neurophysiol 44(4):751–72.

Giszter SF, McIntyre J, Bizzi E (1989) Kinematic strategies and sensorimotor transformations in the wiping movements of frogs. J Neurophysiol 62(3):750–67.

Giszter SF, Mussa-Ivaldi FA, Bizzi E (1993) Convergent force fields organized in the frog’s spinal cord. J Neurosci 13(2):467–91.

Gordon AM, Huxley AF, Julian FJ (1966) The variation in isometric tension with sarcomere length in vertebrate muscle fibres. J Physiol 184(1):170–92

Hart CB, Giszter SF (2004) Modular premotor drives and unit bursts as primitives for frog motor behaviors. J Neurosci 24(22):5269–82.

Hart CB, Giszter SF (2010) A neural basis for motor primitives in the spinal cord. J Neurosci 30(4):1322–36.

Henneman E, Somjen G, Carpenter DO (1965) Excitability and inhibitability of motoneurons of different sizes. J Neurophysiol 28(3):599–620.

Hoffmann P (1910) Beitrag zur Kenntnis der menschlichen Reflexe mit besonderer Berucksichtigung der elektrischen Erscheinungen. Arch Anat Physiol 1:223–246.

Hug F, Avrillon S, Ibáñez J, Farina D. (2023) Common synaptic input, synergies and size principle: Control of spinal motor neurons for movement generation. J Physiol. 601(1):11–20.

Kargo WJ, Giszter SF (2000) Rapid correction of aimed movements by summation of force-field primitives. J Neurosci 20(1):409–26

Kim T, Branner A, Gulati T, Giszter SF (2013). Braided multi-electrode probes: mechanical compliance characteristics and recordings from spinal cords. J Neural Eng10(4)

Kim T, Zhong Y, Giszter SF (2019). Precise Tubular Braid Structures of Ultrafine Microwires as Neural Probes: Significantly Reduced Chronic Immune Response and Greater Local Neural Survival in Rat Cortex. IEEE Trans Neural Syst Rehabil Eng 27(5):846–856.

Kim T, Schmidt K, Deemie C, Wycech J, Liang H, Giszter SF (2019) Highly Flexible Precisely Braided Multielectrode Probes and Combinatorics for Future Neuroprostheses. Front Neurosci 13:613.

Kirkwood PA, Sears TA (1975) Spike triggered averaging for the measurement of single unit conduction velocities. J Physiol 245(2):58–59.

Laine CM, Martinez-Valdes E, Falla D, Mayer F, Farina D (2015) Motor Neuron Pools of Synergistic Thigh Muscles Share Most of Their Synaptic Input. J Neurosci 35(35):12207–16.

Lindén H, Petersen PC, Vestergaard M, Berg RW (2022) Movement is governed by rotational neural dynamics in spinal motor networks. Nature 2022 610(7932):526–531.

Maier MA, Perlmutter SI, Fetz EE (1998) Response Patterns and Force Relations of Monkey Spinal Interneurons During Active Wrist Movement. J Neurophysiol 80(5):2495–2513

McMahon C, Kowalski DP, Krupka AJ, Lemay MA (2022) Single-cell and ensemble activity of lumbar intermediate and ventral horn interneurons in the spinal air-stepping cat. J Neurophysiol 127(1):99–115.

Mitra P, Bokil H (2008) Observed Brain Dynamics. New York City, NY: Oxford University Press.

Negro F, Yavuz UŞ, Farina D (2016)The human motor neuron pools receive a dominant slow-varying common synaptic input. J Physiol 594(19):5491–505.

Paninski L (2004) Maximum likelihood estimation of cascade point-process neural encoding models. Network. 15(4):243–62

Paninski L, Pillow JW, Simoncelli EP (2004) Maximum likelihood estimation of a stochastic integrate-and-fire neural encoding model. Neural Comput. 16(12):2533–61

Poliakov AV, & Schieber MH (1998) Multiple fragment statistical analysis of post-spike effects in spike-triggered averages of rectified EMG. J Neurosci Methods, 79(2), 143–150.

Prochazka A, Gillard D, Bennett DJ (1997) Positive force feedback control of muscles. J Neurophysiol, 77(6), 3226–3236.

Raposo D, Kaufman MT, Churchland AK (2014). A category-free neural population supports evolving demands during decision-making. Nat Neurosci 17(12):1784–1792

McCrea DA, Rybak IA (2008) Organization of mammalian locomotor rhythm and pattern generation. Brain Res Rev. 57(1):134–46.

Sanger TD (2010) Controlling Variability. J Mot Behav 42(6):401–7.

Sanger TD (2011) Distributed control of uncertain systems using superpositions of linear operators. Neural Comput 23(8):1911–34

Sanger TD (2014) Risk-aware control. Neural Comput. 26(12):2669–91.

Schwartz O, Pillow JW, Rust NC, Simoncelli EP (2006) Spike-triggered neural characterization. J Vis. 17:6(4):484–507

Schotland JL, Rymer WZ (1993a) Wipe and flexion reflexes of the frog. I. Kinematics and EMG patterns. J Neurophysiol 69(5):1725–35.

Schotland JL, Rymer WZ (1993b) Wipe and flexion reflexes of the frog. II. Response to perturbations. J Neurophysiol 69(5):1736–48.

Semmler JG, Steege JW, Kornatz WK, Enoka RM (2000) Motor-unit synchronization is not responsible for larger motor-unit forces in old adults. J Neurophysiol 84(1):358–66.

Shiavi R, Negin M (1975) Stochastic properties of motoneuron activity and the effect of muscular length. Biol Cybern 19(4):231–7.

Shoham S, Fellows MR, Normann RA (2003) Robust, automatic spike sorting using mixtures of multivariate t-distributions. J Neurosci Methods 127(2):111–22.

Smith AC, Brown EN (2003) Estimating a state-space model from point process observations. Neural Comput 15(5):965–91

Takei, T, Confais J, Tomatsu S, Oya T, Seiki S (2017) Neural basis for hand muscle synergies in the primate spinal cord. Proc Natl Acad Sci (USA) 114(32):8643–8648.

Thura D, Cabana JF, Feghaly A, Cisek P (2022) Integrated neural dynamics of sensorimotor decisions and actions. PLoS Biol. 20(12)

