## Supplemental Appendix for "A Stochastic Dynamic Operator framework that improves the precision of analysis and prediction relative to the classical spike-triggered average method, extending the toolkit"

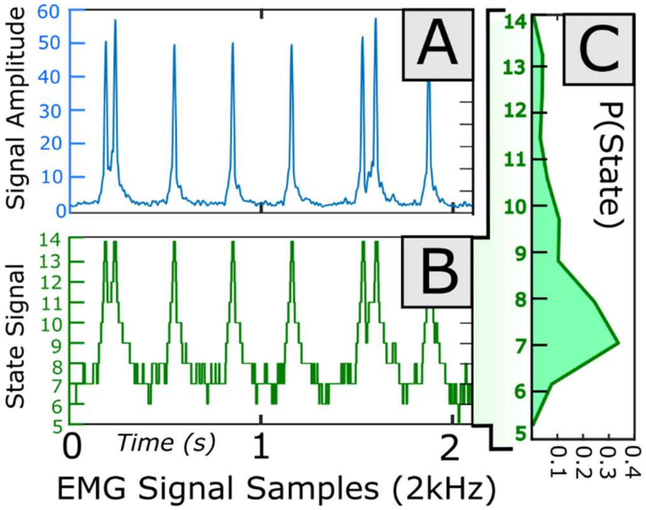


Supplemental Figure 1: Example state quantization of electromyogram (EMG) time series data collected during motor behavior. **A**) Four second sample of rectified and filtered EMG signal. Isolated motor unit responses are evident as peaks in the aggregate EMG. **B**) The ‘stage-signal’ derived from discretizing the EMG time series into binned signal amplitudes. Twenty state bins were defined from logarithmic intervals between the minimum and maximum observed signal amplitude across the dataset (and not observed in the four-second interval shown in A). Hence, the four-second subsample displayed here does not include all potential states of the signal. This state-binning scheme permitted clear segregation between peaks and background, with smooth transitions between subsequent observations (shifting no more than 1 state). **C**) Distributions of signal state may be derived over finite intervals. The plotted distribution of state was drawn from the entire 4 second duration displayed in B. Such distributions can be drawn from arbitrary intervals. Due to these factors, the distribution of signal state may vary by signal dynamics and state binning scheme.

**Signals may demonstrate dynamical behavior irrespective of spike**:

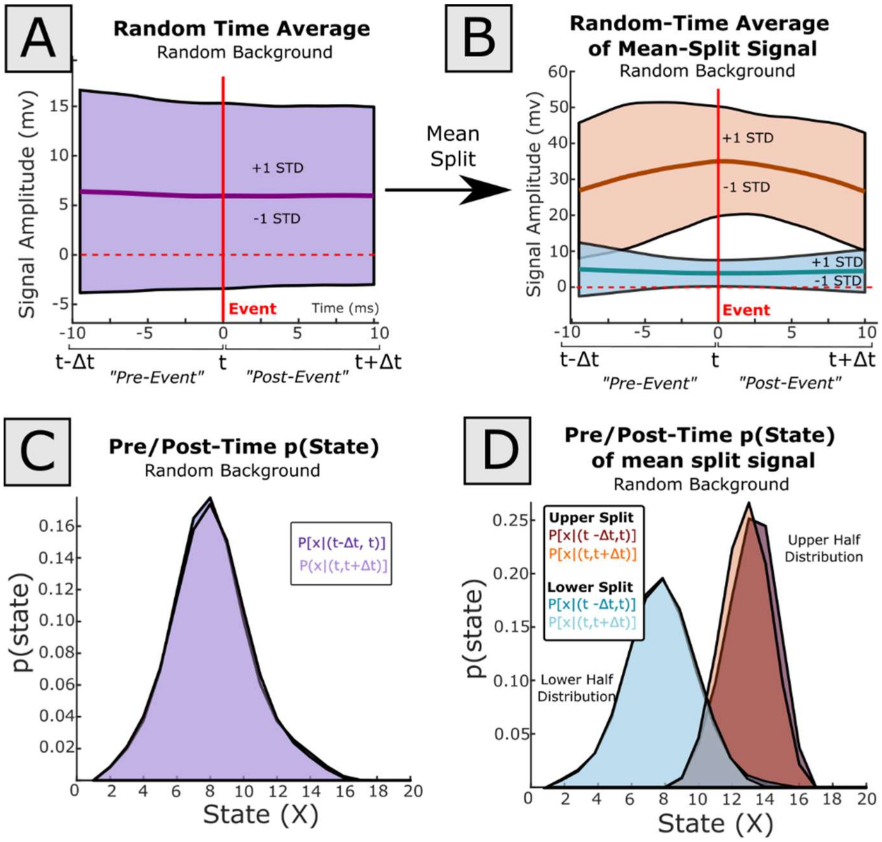


Supplemental Figure 2: The trigger-average of an EMG signal generated from random event times may also vary by state. **A**) The averaged signal waveform shows a flat impulse response, with a broad variance across all event times. **B**) By performing a mean-split on sampled signal waveforms by state at event time, high and low split EMG signal differ in their impulse responses, with a difference in their variance. **C**) Signal amplitude was converted to states using a logarithmic scheme. The average pre-event and post-event time points appear invariant, with equal means and variances. **D**) The mean-split distributions of state pre/post-event differ in mean and variance, indicating that predictions of post-event signal states may be readily predicted from pre-signal state. (Alternatively, signal remains in nearby states after the reference event time, but the distribution of these states may vary by reference state.)

**Computational Details: Ensuring the SDO matrix compliance with theoretical requirements, in data-derived matrices:**

For an SDO to behave as a linear operator manipulating the flow of probability distributions, as described in Sanger, 2010, 2011, there are four conditions which are necessary and sufficient: 1) The matrix diagonal contains non-positive values, 2) The off-diagonal elements contain non-negative values, 3) Each column of the SDO sums to 0, 4) and the sum of all positive elements in a column must be no greater than 1. Multiple matrices could theoretically satisfy the distribution-to-distribution mapping described in EQ2 but may still fail the three conditions necessary to behave as an SDO. Simultaneously, the best linear estimation of the SDO derived from EQ6 is often ill-posed because the covariance matrix of *pre-spike* state distribution ($R_{xx}$ in EQ6) is often non-invertible. To circumvent these problems, we designed an algorithm which will yield a linearly-compliant SDO regardless of input distributions. We describe this process as follows:

State occurrences in the post-spike interval are not independent events: Subsequent states emerge as transitions from prior states. When repeatedly drawing state distributions across an increasing *post-spike* interval, subsequent distributions evolve in time, although perhaps only subtly, reflecting this shared component. To capture this average flow of probability through subsequent states over a defined *pre-spike* and *post-spike* interval, we can average the transitions between every state observation in the *pre-spike* distribution, with every state observation in the *post-spike* distribution. In a doubly-discrete digital signal, as used here, each observation of state in time, *x(t)*, in both the *pre-spike* and *post-spike* intervals can be described as a finite-length binary state probability vector (**Supplemental Figure 3A**).

The outer product of a given pair of post-spike and pre-spike binary state vectors produces a (binary) transition matrix mapping the observed pre-to-post spike state distributions (**Supplemental Figure 3B**). The average of these binary transition matrices, for a given pre-spike state, (i.e., mapped between a single pre-spike binary state probability vector and *all* successive post-spike state binary vectors within the post-spike interval) describes the average state transition, and flow of probability, from this initial state. Accordingly, the *average of the binary transition matrices* mapping the single pre-spike state probability vector to each successive post-spike binary state vector provides the transition matrix between the same pre-spike state and the *average of the post-spike states* (i.e., the *post-spike* state distribution). This process may be repeated for all pre-spike states, generating a transition matrix for each. The average of these pre-spike state transition matrices is thus the overall average state transition between ***all*** *time points* in the pre-spike interval and ***all*** *time points* in the post-spike interval.

Under these assumptions, two transition matrices originating at the same initial state in the pre-spike interval will be identical, because all pre-spike initial states transition to the (same) average *post-spike* state distribution (as described above). Hence, the average transition matrix between the pre-spike states to *post-spike* state distribution depends only on the average frequency of each state in the pre-spike interval, that is, on the *pre-spike* state distribution. Thus, the overall joint distribution of states in the *pre-spike* and *post-spike* distributions can be predicted from the full distributions directly, using an outer-product operation between the two distribution vectors (**Supplemental Figure 3C**). If each column of the joint distribution is normalized to unity, the joint distribution becomes the average (left) transition matrix mapping the *pre-spike* to the *post-spike* distribution in a Markovian framework. (This normalization is necessary for a query probability distribution, premultiplied by the transition matrix, to produce an output probability vector which sums to unity and implies that all output possibilities have been accounted for.)

For every transition between the binary *pre-spike* state distribution vector to the *post-spike* state distribution, there is a (non-binary) vector which captures the differences in these probability distributions (**Supplemental Figure 3D**, left). While the outer product of the *pre-spike* and *post-spike* distribution vectors produces the observed transition matrix, the outer product of the *change* of state vector with the *pre-spike* state distribution produces an array which describes the *change* in the transition matrix. Because the *pre-spike* state distribution within a single bin is binary, the *change* in transition matrix satisfies the SDO linear constraints (above). The average *change in transition* matrix may then be similarly derived as the average of the *change of transition matrices* generated for each pre-spike state observation (**Supplemental Fig 3D**, right). The average *change in transition matrix* obtained in this way is then the linearly-compliant SDO.

The resulting row-sum of the SDO matrix is a column vector which corresponds to the average *change* in the *post-spike* and *pre-spike* state distributions, while each column sums to 0. It is important to note that these two qualities alone are not sufficient to uniquely constrain the resulting *change in transition* matrix. (Indeed, an average *change in transition* matrix may be derived from the outer product of the average *change in state* distribution and initial state distribution.) In most cases, a resulting matrix which only satisfies row and column-sums will not be linearly compliant, and thus will not behave ideally as an SDO. By averaging across binary pre-spike states, our algorithm results in a matrix which maintains these row and column-sums while fully upholding the linearity constraints and is thus compliant. Using this method, the covariance matrix *of pre-spike* distribution ($R_{xx}$ in EQ6) effectively becomes a diagonal matrix of the *pre-spike* state distribution. An alternative derivation relying on these same assumptions can be faster.

***Alternate Direct Method:***

Using the above framework of averaging over binary *pre-spike* state distributions, we can also formulate a more direct (and faster) algorithm for estimating the same SDO matrix derived above. Because the SDO describes the average *change* in the joint distribution of state, and the initial state is first treated as binary, the average transitions from each singular state in the pre-spike interval to the average change in the *post-spike* distribution (the SDO matrix) may instead, and equivalently, be derived from the joint distribution of state, minus the average *pre-spike* state distribution. In the original binary framework, transitions are aggregated from singular pre-spike states to *post-spike* state distributions (**Supplemental Figure 3C**). The equivalent matrix representation of the initial *pre-spike* distribution can be found by averaging the outer product of each component pre-spike **binary** state vector with itself. Because each pre-spike state vector is binary, this operation effectively produces a diagonal matrix, where each diagonal element$_i$ is $prob\big(x,(s-dt,s)\big) = i$. That is, the diagonalized *pre-spike* state distribution. In the alternate (and faster) algorithm, the SDO may be directly derived from the joint distribution of state, less the diagonalized *pre-spike* distribution matrix (**Supplemental Figure 3E**).

This latter algorithm and method of estimation of the SDO thus derives the matrix directly from the observed *pre-spike* and *post-spike* state distributions, then averaged over all spiking events. In addition to parsimony, a second advantage of this algorithm is that both *pre-spike* and *post-spike* state distributions may be filtered and smoothed prior to matrix estimation while retaining the linear compliance of the resulting SDO matrix. Such smoothing may be preferable when the distribution of states cannot be measured precisely (such as over short intervals of discrete signal), or can only be known probabilistically, or in hindsight (e.g., after estimating the probability of event occurrence in time). Towards meeting this end, the *SDO Analysis Toolkit* was written to generate SDOs specifically incorporating this second direct method, with user-defined variable *pre-spike* and *post-spike* interval durations, delays, and smoothing. The first, expositional method, is available as an option in the Toolkit.

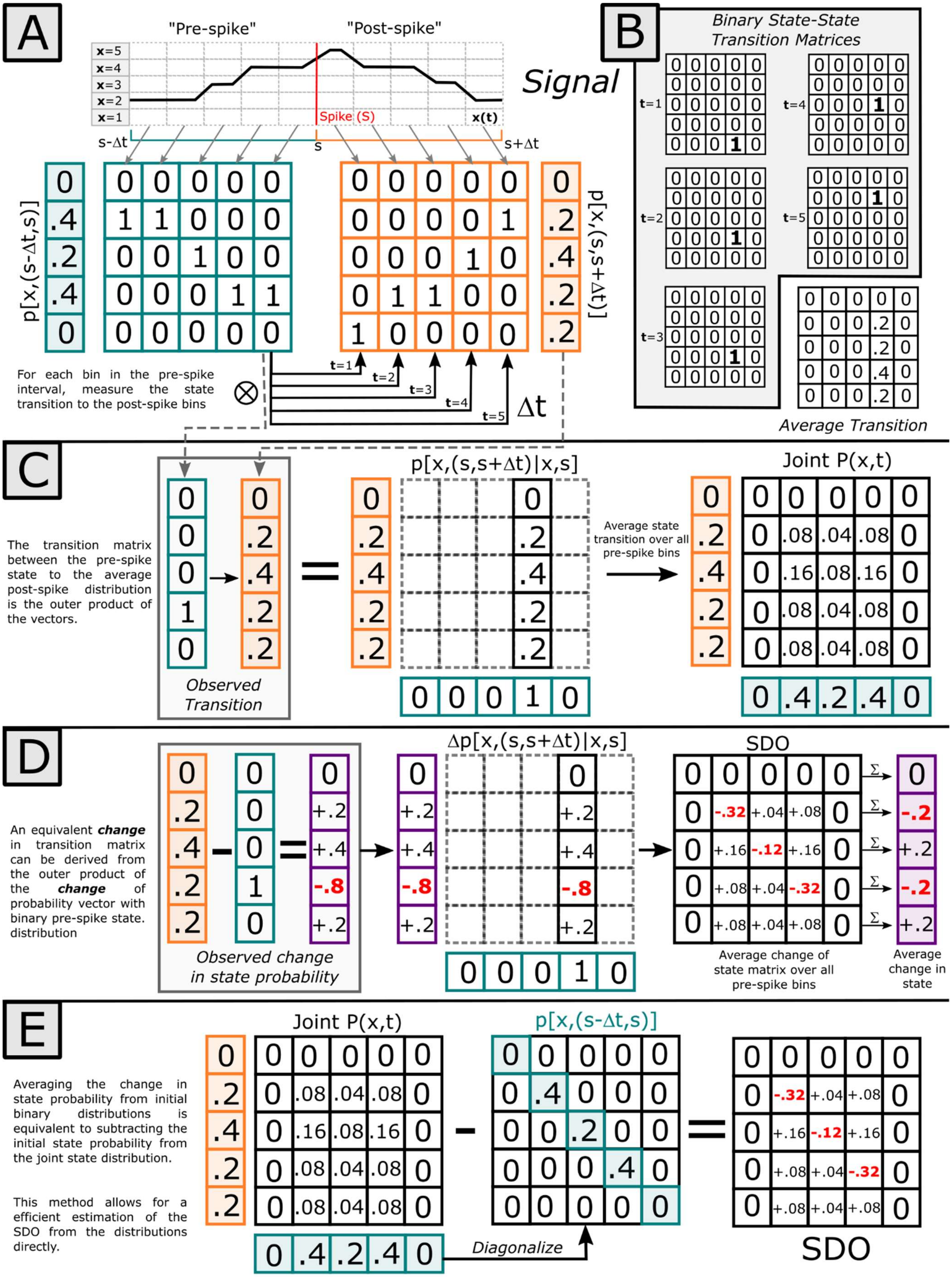
A
"Pre-spike"
"Post-spike"
Signal
x=5
x=4
x=3
x=2
x=1
Spike (S)
x(t)
s-Δt
s
s+Δt
p[x,(s-Δt,s)]
p[x,(s,s+Δt)]
For each bin in the pre-spike interval, measure the state transition to the post-spike bins
t=1
t=2
t=3
t=4
t=5
Δt
B
Binary State-State Transition Matrices
Average Transition
C
The transition matrix between the pre-spike state to the average post-spike distribution is the outer product of the vectors.
Observed Transition
p[x,(s,s+Δt)|x,s]
Average state transition over all pre-spike bins
Joint P(x,t)
D
An equivalent change in transition matrix can be derived from the outer product of the change of probability vector with binary pre-spike state. distribution
Observed change in state probability
Δp[x,(s,s+Δt)|x,s]
SDO
Average change of state matrix over all pre-spike bins
Average change in state
E
Averaging the change in state probability from initial binary distributions is equivalent to subtracting the initial state probability from the joint state distribution.
This method allows for a efficient estimation of the SDO from the distributions directly.
Joint P(x,t)
p[x,(s-Δt,s)]
Diagonalize
SDO

Supplementary Figure 3: Example derivation of the Stochastic Dynamic Operator (SDO) from a single spiking event via the binary vector estimation method. **A**) A digital time series signal, collected in discrete time, can be represented as a sequence of states before and after spike, *s*. Each time bin may be converted to a binary state probability vector, representing the probability of observing state *x* at position *s+t*. The row-wise average across these arrays yields the overall 'pre-spike' or 'post-spike' distributions (Designated as filled boxes). The SDO is the average difference between each bin in the pre-spike interval to each bin in the post-spike distribution. **B**) The transition matrix describing a transformation from a pre-spike to a post-spike state may be derived from the outer product of the post-spike and pre-spike binary state vectors. The average transition between a single pre-spike bin to each post-spike bin is equivalent to the transition between the pre-spike bin and the average of the post-spike bins (i.e., the post-spike distribution). **C**) The joint probability of state in the pre-spike and post-spike bin is thus the averaged transition between each pre-spike bin to the post-spike state distribution (the average of post-spike state). The averaged probability of the pre-spike binary vectors to the average post-spike state probability is equivalent to the transition between the average probability of pre-spike state distribution and the average post-spike state; the joint probability distribution is outer product of the post-spike and pre-spike state distributions. **D**) For each pre-spike binary state vector transition to the post-spike state distribution (average), there is an associated *change in probability* vector (purple). (Negative values are designated in red.) The outer product of this *change of probability* vector with the binary pre-spike state similarly yields a matrix describing the *change* in the joint transition matrix. The average of these change of probability matrices yields the SDO, which describes the overall shift in the joint probability distribution. To preserve the linearity constraints of the SDO (non-positive diagonal elements; non-negative off-diagonal elements; columns sum to 0), this matrix cannot be directly estimated from the outer product of the average *change of probability* vector and the *pre-spike* state distributions. The row sum across this matrix is equal to the average post-spike minus pre-spike distribution vectors. **E**) Algorithmically, and equivalently, the SDO may be equivalently derived as the joint state probability distribution, as derived in **C**, subtracted by the diagonalized pre-spike state probability distribution.

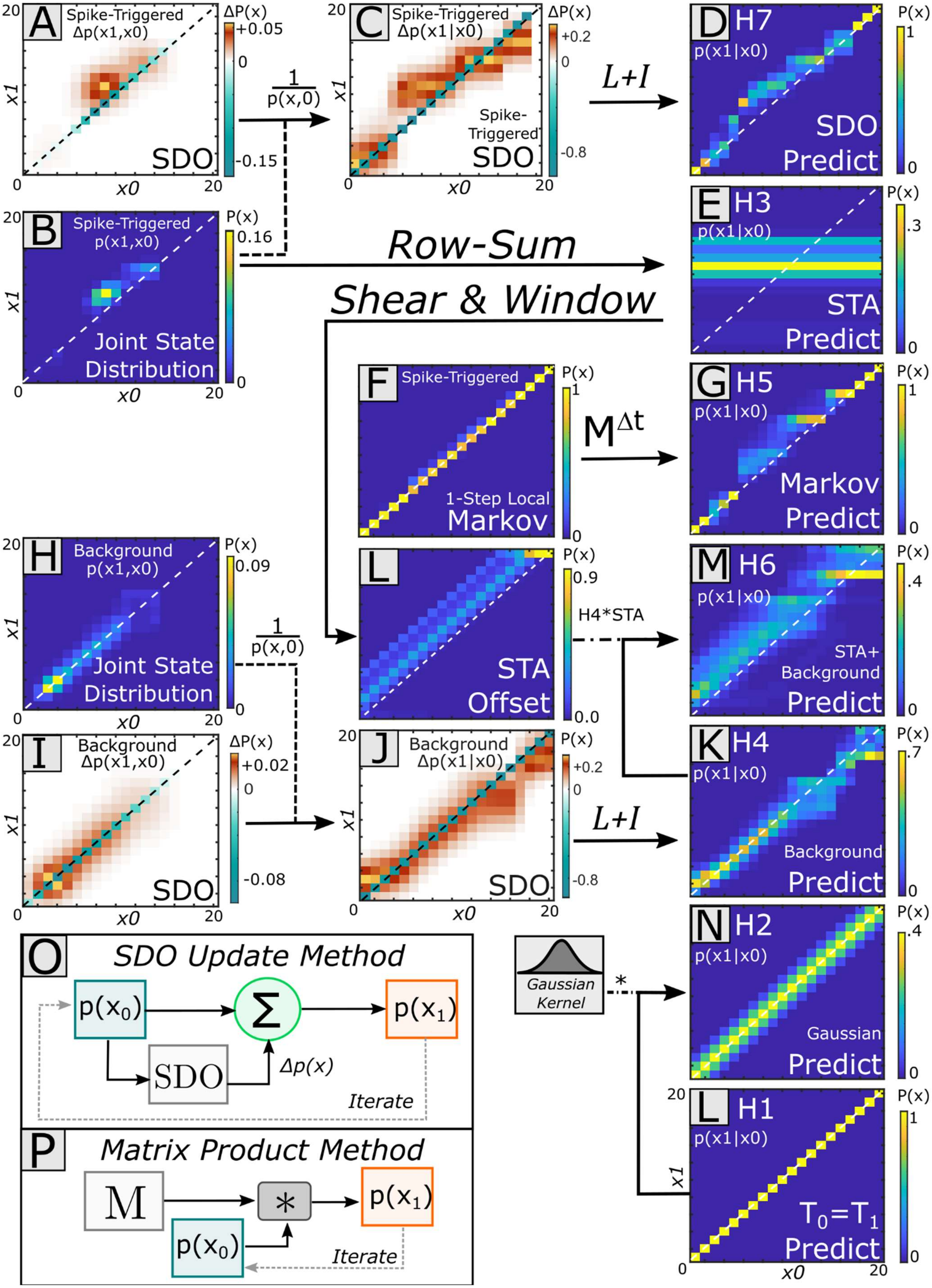

A Spike-Triggered Δp(x1,x0) SDO
B Spike-Triggered p(x1,x0) Joint State Distribution
C Spike-Triggered Δp(x1|x0) Spike-Triggered SDO
D H7 p(x1|x0) SDO Predict
E H3 p(x1|x0) STA Predict
F Spike-Triggered 1-Step Local Markov
G H5 p(x1|x0) Markov Predict
H Background p(x1,x0) Joint State Distribution
I Background Δp(x1,x0) SDO
J Background Δp(x1|x0) SDO
K H4 p(x1|x0) Background Predict
L STA Offset
M H6 p(x1|x0) STA+ Background Predict
N H2 p(x1|x0) Gaussian Predict
L H1 p(x1|x0) T0=T1 Predict
1/p(x,0)
L+I
Row-Sum
Shear & Window
MΔt
H4*STA
Gaussian Kernel
O SDO Update Method
p(x0) Σ p(x1) SDO Δp(x) Iterate
P Matrix Product Method
M * p(x1) p(x0) Iterate

Supplemental Figure 4: Transition Matrix representation of the 7 prediction hypotheses. Pre-spike and post-spike distributions were calculated using 10 ms intervals. The predictions of *post-spike* state distributions Δt steps (1 step = 10 ms) forward in time across all hypotheses are generated from the convolution of the respective transition matrices (**A**). For predictions involving SDOs (H3, H6), the SDO matrix is first normalized to the occurrence of state at spike (p(x)) using the respective joint distribution of state (**B, H**), to produce *change in transition* matrices (i.e., $\Delta p(x_1|x_0)$) (**C, J**). Here, Δt = 1 (i.e., the predicted distribution is '10 ms step' forward; the prediction of the immediate *post-spike* distribution). (Other prediction methods are also possible for the SDO framework, below). **D)** [H7] The spike-triggered SDO captures the spike-triggered dynamics. **E)** [H3] The neuron's (simple) spike-triggered average is derived from the row-sum of the spike-triggered joint distribution of state (**B**). This column vector is then duplicated for all input states. In the resulting transition matrix, the same output distribution is predicted regardless of input distribution. **G**) [H5] A Markov transition matrix is derived from consecutive state transitions observed in the pre-spike interval. This matrix is then convolved over the comparable number of state-transitions in post-spike interval (t0), and further convolved by Δt. **M)** [H6]. Here, the spike-triggered average matrix (H3, **E**) is sheared around the average probability of state. This provides the 'average constant offset of state' for every input state. This sheared matrix (**L**) is then convolved with the background joint distribution of state (**K**) to generate the transition matrix. **K**) [H4] The background SDO is normalized and predicted as with the spike-triggered SDO. **N)** [H2] An identity matrix convolved with a gaussian kernel. This causes an increase in the input *pre-spike* state distribution variance without a shift of mean. **L)** [H1] The identity matrix. The *post-spike* state distribution will resemble the *pre-spike* distribution exactly. There are two methods of predicting the *post-spike* state distribution. **O**) SDO matrices, *L*, predict a *change* of probability, Δp(x), with the predicted output probability as the linear sum of the input state, p(x0) and this change in probability; $p(x_0)+\Delta p(x)=p(x_1)$. This output state distribution can then serve as a new input to iteratively predict forward over an arbitrary interval. **P**) In the transition matrix form, the prediction of *post-spike* state distribution is the transition matrix post-multiplied by the *pre-spike* state distribution.

***Matrix Implementation of 7 Hypotheses***: We generated 7 sets of predicted post-spike distributions from the 7 hypothesized relationships between spike-source and signal (**Supplemental Figure 4**). To generate the predicted *post-spike* state probability distributions at time Δt, we first generated the left transition matrix describing each hypothesis over this interval. When deriving the transition matrix to the joint distribution of state, each column must be normalized to 1 (reflecting that given an input state, the probability of all transitions to the consecutive state must be accounted for). For each column *j,* the joint distribution column must be scaled by a factor $k_j$, such that the column sums to 1; $k_j\ p(x_0=j)=1$. Clearly, this factor $k_j$ is the reciprocal of the joint distribution column, $k_j=\frac{1}{p(x_0=j)}$. Analogously, the background and spike-triggered SDOs require a normalization to properly encode this *change* of state probability. To scale the SDO appropriately, each column *j* is similarly scaled by factor $k_j$ of the matching joint distribution. These transition matrices could then be post-multiplied by an arbitrary *pre-spike* state distribution to generate the predicted *post-spike* distribution.

**<u>[H1] No change</u>** (**Supplemental Fig. 4L**) $\hat{p}(x_1)=p(x_0)$: Here, the transition matrix is the identity matrix; the predicted *pre-spike* distribution is thus identical to the *post-spike* distribution.

**<u>[H2] Gaussian Convolution</u>** (**Supplemental Fig. 4N**) $\hat{p}(x_1) = G * p(x_0)$: Here, the transition matrix is the identity matrix convolved with a gaussian kernel (This exact kernel is user defined within the *SDO Analysis Toolkit*, by default σ=1). This transition matrix corresponds to an increase in the *post-spike* distribution variance, but no change in mean relative to the *pre-spike* distribution.

**<u>[H3] Spike-triggered Average [STA]</u>** (**Supplemental Fig. 4E**) $\hat{p}(x_1) = \bar{P}(x_1)$: The spike-triggered average is pre-spike state invariant: For all input states, the predicted output distribution is the same. Hence, represented as a transition matrix, each column of the matrix is identical: The encoded distribution of state is the average *post-spike* distribution. This distribution will be recovered regardless of *pre-spike* state distribution. The row-sum across the spike-triggered joint distribution of state is equal to the average *post-spike* state distribution.

**<u>[H4] Background Dynamics</u>** (**Supplemental Fig. 4K**) $\hat{p}(x_1) = L_B p(x_0) + p(x_0)$: The background SDO, calculated for all time points within the signal, was converted into at transition matrix (using the above normalization). This operator is designed to capture the inherent dynamics in signal behavior not correlated with spikes (e.g., regression of signal to a baseline).

**[H5] Local (non-spike associated) probability dynamics (Supplemental Fig. 4G)** $\hat{p}(x_1) = M_0(x_0)$: This is a (left) Markov transition matrix, estimated from sequential observations of state, only in the pre-spike interval. The 1-Step Markov transition matrix is then convolved over the duration of the pre-spike interval.

**<u>[H6] Dynamic Background + Spike-Triggered Average [STA]</u>** (**Supplemental Fig. 4M**) $\hat{p}(x_1) = (L_B)p(x_0) + p(x_0) + \Delta\bar{P}(x)$: This hypothesis tested the combination of state-dependent, non-spike associated background dynamics, with a constant (pre-spike state invariant) offset associated with spike (averaged spike effect). The STA-transition matrix (H3) was sheared relative to the average (overall) *pre-spike* state to produce a diagonal-banded matrix, then column-normalized to 1 (i.e., the constant offset matrix represents the <u>average</u> *post-spike* state distribution, given the <u>average</u> pre-spike state, now applied to all states). This sheared matrix is then convolved with the dynamic background matrix (H4). H6 thus resembles H4 with a constant shift of mean and variance, provided by H3.

**<u>[H7] Dynamic Background + Dynamic Spike Effects [SDO]</u>** (**Supplemental Fig. 4D**) $\hat{p}(x_1) = Lp(x_0) + p(x_0)$: The spike-triggered SDO, calculated from the transitions between the *pre-spike* and *post-spike* distributions was converted into a transition matrix (using the above normalization).

**Extended Details on Significance Tests of SDO Matrices**:

To identify significant SDOs within our dataset, we explored four measures of significant effect and one measure of state-tuning. For each measure, the spike-triggered SDO test statistic was compared against the distribution of test statistics generated from the 'shuffled' SDOs using the shuffled spike trains. Here, the distance between the spike-triggered SDO measure and the average 'shuffled' SDO was compared to the distance between each 'shuffled' SDO and the mean of the shuffles (i.e., internal variance). A spike-triggered SDO test statistic was considered 'significant' if the distance between the spike-triggered SDO and the mean of the shuffle was in the top 5% of observed effects (i.e., a 1-tailed effect, $\alpha = 0.05$). The number of shuffles to draw and the alpha value for significance may be adjusted within the *SDO Analysis Toolkit* as desired. The 'shuffled' SDOs were reparameterized to reflect the same *pre-spike* distribution of state as the spike-triggered SDO prior to taking significance measures. These measures are as follows:

1. <u>The element-wise difference in magnitude (finding Significant Elements)</u>: For each element of the SDO matrix the sum of squared distance (SSD) was calculated between the element of the spike-triggered SDO and the mean of matching element of the shuffled-spike SDOs. This distance was compared, on an elementwise basis, against the distribution of distances calculated between each (1000) shuffled-spike SDO and the mean of the shuffled-spike SDOs. The *SDO Analysis Toolkit* includes the option to z-score normalize these values prior to significance testing. SDO elements were considered significant if their distance to the mean value was >95$^{th}$ percentile, relative to shuffled spikes. This threshold was Bonferroni-corrected for the number of independent tests (i.e., the number of elements; 20x20 = 400).
2. <u>The sum of element-wise SSD in magnitude (testing for matrix Significance)</u>:The sum of the squared differences between the spike-triggered and mean shuffled SDOs were accumulated (summed) over all elements $\sum_i \sum_j \left|SDO_{i,j} - Shuff_{i,j}\right|^2$. This cumulative SSD was similarly calculated between each shuffled SDO and the mean of the shuffled SDOs. Spike-triggered SDO matrices were considered broadly significant if the cumulative SSD was >95$^{th}$ percentile of the null/shuffled SDO effect.
3. <u>State-wise Coarse output state bias (directional shifts towards higher or lower states)</u>: For a given input state, the spike-triggered SDO may, on average, increase, decrease, or sustain the mean state value in

the predicted *post-spike* distribution. Directional effects of the mean predicted *post-spike* state distributions from the SDO were assessed for every input state. This directional effect was calculated as the *difference* between 1) sum of column elements above the diagonal, (corresponding to *post-spike* states > *pre-spike* states) and 2) the sum of column elements below the matrix diagonal (corresponding to *post-spike* states < *pre-spike* states). This value corresponds to the net shift in the mean of *post-spike* state probability distribution, given an input pre-spike state. (Because increased signal amplitudes are assigned higher states, a coarse bias greater than zero indicates a transition of the mean probability differences towards higher states, negative towards lower states, and 0 of sustaining the mean of the *post-spike* distribution at the input state.) This measurement is reflected in the 'Quiver' visualization of the SDO. This threshold was Bonferroni-corrected for the number of states used (20).

4. Array-wise Cumulative Coarse state basis (directional shifts towards higher or lower states): Column-wise input state bias was calculated independently for all states, as above, and concatenated into a row vector for all input pre-spike states. Here, the overall array effect was tested, that is, calculating the magnitude overall bias of the SDO matrix, relative to biases of shuffled-spike elements. (This permits accumulated effects to differ, even if individual state-wise coarse biases do not reach threshold. E.g., A significant SDO may contain a higher frequency of state-wise directional effects than the null, even if the magnitude of each effect in isolation is not significantly different from the null distribution.) We calculated the SSD between this coarse state bias vector and the equivalent coarse state bias vector calculated from the mean of the shuffled-SDO. The SSD distance between the coarse state bias between each shuffled-SDO and the mean-shuffled SDO was similarly calculated. The cumulative spike-triggered coarse state bias was considered significant if the spike-triggered SSD value was in the >95th percentile relative to the shuffled-SDO distribution.
5. The probability of state at spike (State tuning): The distribution of states occurring immediately prior to spike (*pre-spike* distribution) could differ from background distribution of signal state. To determine significant effects, the Kullback-Leibler Divergence (KLD) was calculated between the average *pre-spike* distribution to the mean of the 'shuffled' *pre-spike* distributions. A distribution of KLD values was calculated similarly between each shuffle and the mean of the shuffles. The spike-triggered SDO was

considered to have significantly different *pre-spike* state distributions if the observed KLD value was >95th percentile of the shuffled KLD distribution. (By definition, this can only be measured on the original, non-reparameterized, SDO matrices). This measure is thus not a measure of the SDO 'effect', but rather recruitment of the spiking event by signal state.

**Assessing Prediction Accuracy of SDO Matrix Hypotheses**:

Predicting Distributions: The similarities between the ground truth (observed) *post-spike distribution* and the 7 hypothesis-predicted *post-spike* signal state distributions were assessed using both the KLD and the log-likelihood, calculated independently for all spiking events. The log-likelihood of the experimental *post-spike* state distribution was calculated against itself to provide a theoretical limit on maximum likelihood. Because the distribution of KLD and log-likelihood values collected over all spiking events was not gaussian, we used the median as the measure of central tendency for each hypothesis. A bootstrapping technique was used to calculate the 95% confidence intervals of the median to identify significant differences.

Predicting Most Likely Single States: For each spiking event, we assigned the mode of the *post-spike* distribution, as generated by each of the six hypotheses, as the most likely single state. Each spike, and associated errors, was treated as an independent observation. We describe an $e_0$, $e_1$, and $e_2$ error to quantify the differences between predicted and observed post-spike states. The $e_0$ error is the frequency of error, the total number of incorrect predictions. The $e_1$ error is the mean absolute error, the magnitude of the difference between the observed and post-spike state, summed over all spikes. The $e_2$ error is the sum of the squared differences (SSD) between the observed and predicted values, summed over all spiking events.

These prediction metrics are scalar quantities, generated on a spike-wise basis, and are not expected to have a zero-mean or to fall in a normal distribution. To use a common, but robust, method to determine the significance of our predictions, we used a bootstrapping technique. For each prediction metric, we generated 1000 shuffles of the spike-wise error associated with each hypothesis, using the same number of spikes as the original spike train. We summed over the errors in each shuffle to create a distribution of the cumulative error metric associated with each hypothesis (i.e., bootstrapping the cumulative error). We used 95% Confidence intervals for classifying significant differences between bootstrapped distributions. Significant differences in $e_1$ or

$e_2$ errors between groups were assessed by multiple-comparisons tests with Bonferroni correction for the number of unique comparisons. We visualized and analyzed prediction data using our *SDO Analysis Toolkit*, written in MATLAB, which utilizes standard tools from the *Statistics and Machine Learning* Toolbox.

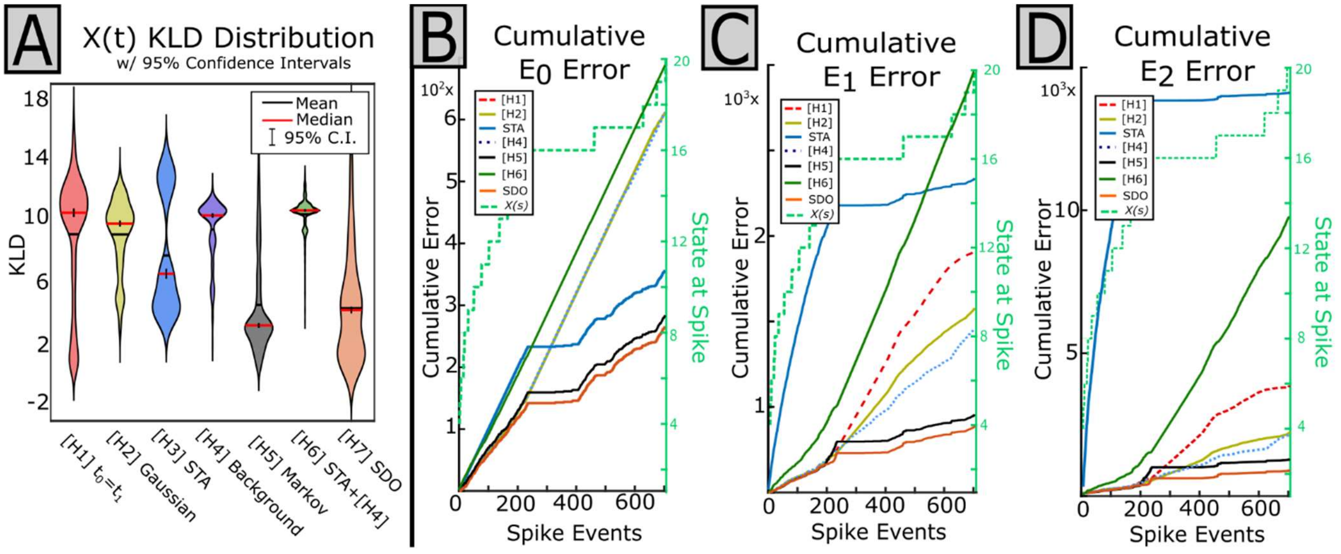


Supplemental Figure 5: Extended prediction statistics, demonstrating all 7 prediction hypotheses for an interneuron vs. the aggregate muscle electromyogram in the vastus externus. **A**) The distribution of KLD statistics recovered between the predicted distributions generated by each of the 7 hypotheses against the observed distributions of state. Here, a lower KLD statistic indicates a closer fit between the predicted and observed distributions. Overall, the SDO has the lowest average prediction error. **B)** The cumulative frequency (E0), **C)** magnitude (E1) and **D)** squared magnitude (E2) error for the 7 hypotheses. The SDO and STA both improve predictions relative to the background dynamics. When predicting to single states in this instance, the SDO produces the lowest prediction error. Here, the SDO and the 1st-order Markov behave similarly. Including the background distribution into the STA [H6] reduces prediction error, but still underperforms the SDO.

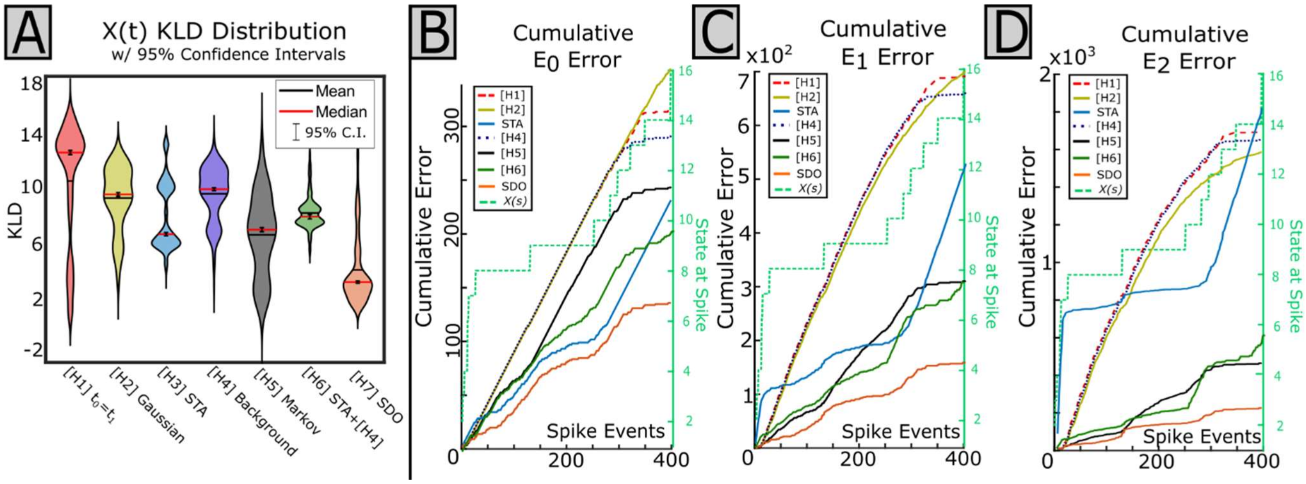


Supplemental Figure 6: Extended prediction statistics, demonstrating all 7 prediction hypotheses for a single motor unit in the vastus externus against the aggregate EMG activity in the biceps femoris. **A**) The distribution of KLD statistics recovered between the predicted distributions generated by each of the 7 hypotheses against the observed distributions of state. Here, a lower KLD statistic indicates a closer fit between the predicted and observed distributions. Overall, the SDO has the lowest average prediction error. **B**) The cumulative frequency (E0), **C)** magnitude (E1) and **D)** squared magnitude (E2) error for the 7 hypotheses. When predicting to single states in this instance, the SDO produces the lowest prediction error. This improved performance is particularly useful when predicting from a higher state (> 10).

**Supplemental References**:

Sanger TD (2010) Controlling Variability. J Mot Behav 42(6):401-7.

Sanger TD (2011) Distributed control of uncertain systems using superpositions of linear operators. Neural Comput 23(8):1911-34
